# Taxonomic difference in marine bloom-forming phytoplanktonic species affects dynamics of both bloom-responding prokaryotes and prokaryotic viruses

**DOI:** 10.1101/2023.01.31.526402

**Authors:** Hiroaki Takebe, Kento Tominaga, Tatsuhiro Isozaki, Tetsuhiro Watanabe, Keigo Yamamoto, Ryoma Kamikawa, Takashi Yoshida

## Abstract

The production of dissolved organic matter during phytoplankton blooms and consumption by heterotrophic prokaryotes promotes marine carbon biogeochemical cycling. Although prokaryotic viruses are crucial biological entities, their dynamics during such blooms are not fully understood. Here, we investigated the dynamics of coastal prokaryotic communities and viruses during blooms in a microcosm experiment using dissolved intracellular fractions of taxonomically distinct phytoplankton, the diatom *Chaetoceros* sp. (CIF) and the raphidophycean alga *Heterosigma akashiwo* (HIF). Ribosomal RNA gene amplicon and viral metagenomic analyses revealed that particular prokaryotes and prokaryotic viruses specifically increased in either CIF and HIF, indicating that different phytoplankton intracellular fractions promote distinct dynamics of not only prokaryotic community but also prokaryotic viruses. Our microcosm experiments and environmental data mining identified both known and novel possible host-virus pairs. In particular, a growth of phytoplanktonic organic matter-associated prokaryotes, such as Bacteroidetes *Polaribacter* and NS9 marine group, *Vibrio* spp., and Rhodobacteriales *Nereida* and *Planktomarina*, was accompanied by an increase in viruses predicted to infect Bacteroidetes, *Vibrio*, and Rhodobacteriales, respectively. Collectively, our findings suggest that elucidating tripartite relationships among phytoplankton, prokaryotes, and prokaryotic viruses would further our understanding of coastal microbial ecosystems.

We state that

-All the data underlying the study are available at the DNA Data Bank of Japan (DDBJ) under project number PRJDB14359 and accession number DRA014887.

-This study was supported by Grants-in-Aid for Scientific Research (No. 16H06429, No. 17H03850, No. 21H05057, and No. 21J14854) from the Japan Society for the Promotion of Science (JSPS).

-We have no potential conflicts of interest to declare.

-We have read and understood your journal’s policies, and we believe that neither the manuscript nor the study violates any of these.

-We do not use any clinical data, human subjects, or laboratory animals.

-None of the materials have been published or are under consideration for publication elsewhere.

**CRediT:** Hiroaki Takebe: Funding acquisition, conceptualization, investigation, formal analysis, visualization, and writing (original draft). Kento Tominaga: Conceptualization, investigation, and writing (review and editing). Tatsuhiro Isozaki: investigation. Tetsuhiro Watanabe: Resources. Keigo Yamamoto: Resources. Ryoma Kamikawa: Conceptualization, supervision, and writing (review and editing). Takashi Yoshida: Funding acquisition, project administration, conceptualization, supervision, and writing (review and editing).

## Introduction

Marine dissolved organic matter (DOM) is one of the major carbon pools on Earth (Hedges, 1992; Hansell and Carlson, 2001). Phytoplankton are mainly responsible for marine primary production, which is known to be comparable to terrestrial levels, and are the major source of marine DOM (Moran *et al*., 2016). Some species like *Heterosigma akashiwo* (Raphidophyceae) and *Chaetoceros* spp. (Diatomea) eventually form blooms in coastal areas through local and seasonal massive growth (Buchan *et al*., 2014; Needham and Fuhrman, 2016; Tomaru *et al*., 2018; Matcher *et al*., 2021). Phytoplankton blooms are often constituted by multiple species (Buchan *et al*., 2014; Nowinski *et al*., 2019) and organic matter produced by each varies species (Brown, 1991; Sarmento *et al*., 2013; Tada and Suzuki, 2016; Tada *et al*., 2017). Thus, DOM released during blooms comprises complex mixtures of a vast range of different organic molecules (Buchan *et al*., 2014).

A large portion (10–50%) of the DOM pools produced and released by phytoplankton is consumed and converted into smaller molecules by heterotrophic prokaryotes (Azam *et al*., 1983). Recent metagenomic and metaproteomic studies in natural blooms have revealed that the relative abundance of specific prokaryotic taxa, such as Flavobacteriales (Bacteroidia), Rhodobacteriales (Alphaproteobacteria), Alteromonadales (Gammaproteobacteria), and Oceanospirales (Gammaproteobacteria), which are capable of utilizing DOM released from phytoplankton, increased drastically within a week (Teeling *et al*., 2012, 2016; Buchan *et al*., 2014; Needham and Fuhrman, 2016). Based on microcosm experiments, patterns of shifts in taxonomic composition and diversity of prokaryotic communities were suggested to depend on bloom-forming phytoplankton species (Landa *et al*., 2014; Tada and Suzuki, 2016; Tada *et al*., 2017). This is explained by the fact that DOM produced varies among phytoplankton species and utilizable and preferred organic molecules vary among prokaryotes (Alonso-Sáez and Gasol, 2007; Sarmento *et al*., 2013). One of those studies further reported that shifts in the prokaryotic community led to changes in amino acid composition (Tada *et al*., 2017), indicating that the DOM is actually consumed and converted by heterotrophic prokaryotes. Accordingly, the phytoplanktonic composition in blooms affects the heterotrophic prokaryote community structure and the resultant biogeochemical cycles (Buchan *et al*., 2014).

In addition to their relationships with phytoplankton, the ecology of these heterotrophic prokaryotes and the resultant biogeochemical cycles are affected by other biotic interactions in the natural environment (Fuhrman, 2009; Martinez-Garcia *et al*., 2012; Needham *et al*., 2013). In particular, viruses are the most abundant biological entities and many infect marine prokaryotes (Suttle, 2005, 2007; Breitbart *et al*., 2018). A theorical study estimated that viral infection to prokaryotes contributes to 10–20% of prokaryotic cell mortality per day (Suttle, 1994) and a most recent study analyzing prokaryotic and viral dynamics in a coastal seawater reported that most of abundant prokaryotic taxa are frequently infected in the natural environment (Tominaga *et al*., in press). Those viral infection enable the release of their cellular compounds and metabolites into DOM pools, which can be readily taken up by other microorganisms (Breitbart *et al*., 2018; Suttle, 2005, 2007). Some prokaryotic viruses also modulate host metabolism for efficient infection through genes called viral auxiliary metabolic genes (AMGs), which are evolutionarily derived from their host genomes (Rosenwasser *et al*., 2016; Zimmerman *et al*., 2020). Therefore, to understand microbial ecology involving biogeochemical cycles in natural blooms formed by multiple phytoplankton species, their effect on the dynamics of prokaryotic viruses also needs to be elucidated. A previous study conducting microscopic observations of viral particles suggested that viral production is active during natural phytoplankton blooms (Matteson *et al*., 2012). Furthermore, a recent study combining isolation techniques and metagenomic analysis found that taxonomically diverse Flavobacteriales viruses emerged during a natural bloom of mostly diatoms (Bartlau *et al*., 2021). However, the ecological and taxonomic links between bloom-forming phytoplankton, marine heterotrophic prokaryotes, and prokaryotic viruses remain unclear.

In the present study, we aimed to observe the effects of DOM released from phytoplankton on prokaryotic community shifts and dynamics of prokaryotic viruses. We performed a microcosm experiment in which the dynamics of coastal prokaryotic and viral communities were analyzed by 16S rRNA gene amplicon and viral metagenomic analyses, respectively, during incubation under laboratory conditions with intracellular fractions obtained from cultured strains of globally distributed bloom-forming species: *H. akashiwo* (Chang *et al*., 1990; Nagasaki *et al*., 1994; Needham and Fuhrman, 2016; Matcher *et al*., 2021) and *Chaetoceros* sp. (Booth *et al*., 2002; Begum *et al*., 2015; Needham and Fuhrman, 2016; Tomaru *et al*., 2018). Furthermore, we analyzed the environmental metagenomic data obtained from Osaka Bay, Japan (Tominaga *et al*., in press), to confirm whether the prokaryotic and viral dynamics observed in our experiment occurred in the natural environment. We propose that different phytoplankton blooms lead to different dynamics of prokaryotic viruses and identified several possible host-virus pairs associating the specific bloom-forming species that have not been reported.

## Experimental Procedures

### Culturing of phytoplankton and collection of intracellular fractions

*H. akashiwo* strain NIES-293 and *Chaetoceros* sp. strain NIES-3717 were purchased from the National Institute for Environmental Studies (NIES, Tsukuba, Japan) and grown axenically in 300 mL of f/2 medium (Guillard and Ryther, 1962) at 20 °C under 10/14 h light/dark photocycle conditions at 40 µmol photons m^-2^ s^-1^. Phytoplanktonic cells in the exponential growth phase were collected by centrifugation at 420 × *g* for 10 min (High Capacity Bench-top Centrifuge LC-220, TOMY SEIKO, Tokyo, Japan). The cell pellet was washed twice with autoclaved aged seawater and resuspended in 30 mL of the same. The washed cells were fractured using a bead-beater (µT-12, TAITEC, Saitama, Japan) (3,200 rpm, 60 s ×2) with 1 cm^3^ of glass beads (0.5 mm diameter, AS ONE, Osaka, Japan), which had been pre-combusted at 200 °C for 2 h. Crushed cellular debris including membranous fractions and glass beads were removed by centrifugation (13,000 × *g* for 15 min), and contaminated glass beads in the supernatant were subsequently removed by filtration using a polyvinylidene cellulose acetate filter (25 mm diameter, 0.2 μm pore size, Millipore, Billerica, MA) to obtain 30 mL of dissolved intracellular fractions including organic matter. The dissolved intracellular fractions derived from *Chaetoceros* sp. and *H. akashiwo* were named CIF and HIF, respectively. CIF and HIF were stored at −80 °C until use. To measure the carbon concentration in the CIF and HIF, 1.5 mL of each sample was diluted with 10 times the volume of Milli-Q water and pre-filtered with 25 mm diameter, 0.2 μm pore size filters (Millipore). The total organic carbon concentration of these samples was analyzed using a total organic carbon analyzer (TOC-L, Shimadzu, Kyoto, Japan).

### Experimental setup

Approximately 12 L of seawater was collected at 3.8 m depth of Osaka Bay, Japan (34°19’28″ N, 135°7’15″ E), on January 31, 2019. The seawater was pre-filtered through polycarbonate membrane filters (142 mm diameter, 3.0 μm pore size; Millipore) to remove eukaryotic cells. For the microcosm experiments, 1.2 L of the pre-filtered seawater was filtered through polycarbonate membrane filters (142 mm diameter, 0.2 μm pore size; Millipore) to obtain prokaryotic cells and reduce the floating viral particles. The prokaryotes and viruses on the filters were re-suspended in 1.2 L of autoclaved seawater in 2 L flasks. Residual organic matter on the surface of the flasks was removed by washing with 6 N HCl followed by Milli-Q water before use. Five milliliters of CIF or 4.43 mL of HIF was added to the seawater samples containing the prokaryotic fraction (84 µmol C/L final concentration). The input carbon concentration was adjusted to increase that caused by natural phytoplankton blooms (Børsheim *et al*., 1999). Control, CIF-treated and HIF-treated flasks were collectively called as control, CIF-treatment and HIF-treatment, respectively. Triplicate flasks of each treatment were designated as replicate flasks I-III. Nine flasks in total were incubated at 20 °C under a 14.5/9.5 h light/dark photocycle condition at 150 µmol photons m^-2^ s^-1^ for 7 days. Temperature and light/dark conditions were consistent with the representative environmental conditions of the bloom season in Osaka Bay, Japan.

### Counting of prokaryotic cells and viral particles

Incubated control, CIF-treatment, and HIF-treatment flasks were homogenized by mixing once a day before subsampling. For counting prokaryotic cells and viral particles, those in 1,920 µL were subsampled from each flask every day and were added to glutaraldehyde (1% final concentration) for fixation at 4 °C. The fixed prokaryotic cells in half of each fixed sample were stained with SYBR® Green I (final concentration 1×; Thermo Fisher Science, Waltham, MA, USA) for 20 min at room temperature under dark conditions. For viral counting, prokaryotic cells in the remaining aliquots of fixed samples were removed by filtration with a cellulose acetate filter (13 mm diameter, 0.2 μm pore size; ADVANTEC Toyo Kaisha, Ltd., Tokyo, Japan). The filtrates were subjected to staining with SYBR® Green I (final concentration 1×) for 20 min at 80 °C, followed by incubation at room temperature according to a previous study (Brussaard, 2004). At least 10,000 stained cells and viral particles were counted using an S3e Cell Sorter (Bio-Rad, Hercules, CA, USA) and analyzed using FlowJo (Becton, Dickinson and Company, Franklin Lakes, NJ, USA), according to manufacturer’s instructions. The CIF-treatment flask replicate III sample on day 3 was lost owing to a technical error. Mann–Whitney U tests were performed to compare the cell and viral abundance of samples in the same phase (early, middle, late) among the control, CIF-treatment, and HIF-treatment flasks.

### DNA extraction and sequencing

For prokaryotic community structure analysis, 1,800 µL of the sample was subsampled from each flask every day (72 samples in total) and the cells collected on polycarbonate membrane filters (13 mm diameter, 0.2 μm pore size; ADVANTEC Toyo Kaisha, Ltd.) and stored at −30 °C until DNA extraction. For community composition in original seawater, prokaryotic cells in 100 mL of seawater were collected on polycarbonate filters (47 mm diameter, 0.2 μm pore size; ADVANTEC Toyo Kaisha, Ltd.), and the filters were stored at −30 °C until DNA extraction. DNA was extracted using previously published methods (Takebe *et al*., 2020). The 16S rRNA genes were amplified using a primer set for the V3–V4 hypervariable region of prokaryotic 16S rRNA genes (Takahashi *et al*., 2014) with added overhang adapter sequences at each 5ʹ-end, according to the manufacturer’s sample preparation guide (https://support.illumina.com/documents/documentation/chemistry_documentation/16s/16s-metagenomic-library-prep-guide-15044223-b.pdf). The amplicons were sequenced using the MiSeq Reagent Kit, version 3 (2×300 bp read length; Illumina, San Diego, CA, USA).

For viral community structure analysis, 100 mL of the sample was subsampled from each flask on days 0, 1, 3, 5, and 7 (45 samples in total). To analyze the viral composition in the original seawater, 100 mL of 3.0 μm pre-filtered seawater was stored. Prokaryotic cells were removed from the 46 samples by filtration with 0.22 μm pore Sterivex filtration units (SVGV010RS, Millipore) and filtrates were stored as viral fractions at 4 °C until DNA extraction. The viruses in the filtrate were concentrated via FeCl_3_ precipitation and purified by CsCl density centrifugation, followed by DNase treatment as described in a previous study (Hurwitz *et al*., 2013). DNA extraction was performed using the combination of xanthogenate-SDS and PCI/CIA protocols described in a previous study (Kimura *et al*., 2012). Extracted DNA was stored at −30°C. Libraries were prepared using a Nextera XT DNA sample preparation kit (Illumina) according to the manufacturer’s protocol, except for the amount of used DNA; we used 0.25 ng of viral DNA as input while manufacturer’s recommendation was 1 ng of DNA. To obtain reference viral genomes in the microcosm experiments, we sequenced reads from one flask in which we observed the most abundant viral particles for each treatment. The remaining samples (two flasks per treatment) were sequenced for mapping onto the reference genomes to analyze the dynamics of viruses. Libraries for two samples (control replicate II on days 1 and 3) could not be prepared; thus, we removed them for the following procedure. The retained libraries were sequenced using MiSeq V3 (2 × 300 bp) (Illumina).

### Sequence processing and ASV generation

From the original seawater sample, 25,220 reads were obtained (Supporting Information Table S1). During the microcosm experiments, 345,382 reads in the control and 308,474, and 683,932 in the CIF- and HIF-treatment flasks, respectively, were obtained on average per flask (Supporting Information Table S1). The DADA2 plugin (Callahan *et al*., 2016) of QIIME2 was used for quality filtering, denoising, pair-end merging, and construction of a feature table of amplicon sequence variants (ASVs). Taxonomic assignment of ASVs was performed using the SILVA (release 138) reference database (Quast *et al*., 2013) and those assigned to mitochondrial or chloroplast sequences were removed from the feature table for further analyses. The singleton ASVs were removed during this step and the control flask replicate I sample on day 1 was removed from further analyses as the obtained reads were less than 5000.

### Analysis of community structure and dynamics of prokaryote ASVs

Rarefaction curves were constructed for all samples using the “iNEXT” package in R (Hsieh *et al*., 2016). Sequence depths were likely sufficient for all samples as the ASVs in those samples were almost saturated by 10,000 reads (Supporting Information Fig. S1); thus, we retained all the samples, except for control flask replicate I on day 1 (see above) for further analyses. For ASV composition, all sequence data were rarefied to the lowest sample depth of 9,450 reads per sample, the Bray–Curtis dissimilarity index calculated using “vegan” in R for pairwise comparisons of the prokaryotic communities in all samples, and then visualized by principal coordinate analysis (PCoA) using the “stats” package in R. Analysis of similarity (ANOSIM) was performed with the Bray–Curtis dissimilarity scores to assess significance of the differences between early (day 0–1) and middle-late (day 2–7) samples and that among middle-late samples between the control, CIF- and HIF- treatments using the R package “vegan.” *P*-values for multiple comparisons were corrected using the Bonferroni method.

The relative abundance of each ASV was calculated by dividing the read count of the ASV by the total read count of each sample. An approximate cell number (ACN) of each ASV was obtained multiplying the relative abundance (0–1) by the total number of prokaryotic cells. Certain ASVs were regarded as abundant if they ranked in the top 20 in ACN at least one day after day 1 in all triplicate flasks and their ACN became more than twice as high as day 0 at least once. We used LEfSe (Segata *et al*., 2011) to test whether differences in the ACN of ASVs of interest among treatments were statistically significant, using treatment as a class and triplicate flasks as subclasses.

### Survey of prokaryote ASVs abundance in amplicon dataset from Osaka Bay natural seawater samples

To confirm that ASVs of interest were detected in the natural environment, we analyzed their abundance at our original sampling site, Osaka Bay. Raw amplicon reads of 16S rRNA gene sequences obtained from natural seawater samples collected monthly in Osaka Bay between May, 2015, and November, 2016 (Tominaga *et al*., in press), were retrieved from the DNA Data Bank of Japan (DDBJ) under project number PRJDB10879. These reads were quality-trimmed and merged, and chimeras were removed following the pipeline described in a previous study (Takebe *et al*., 2020). Quality-controlled reads were mapped on ASVs of our interest with 100% identity using VSEARCH (Rognes *et al*., 2016).

### Sequence processing, vOTU clustering and taxonomic classification

We obtained 17,171,182–25,143,457 reads and 1,159,097–3,806,753 viral metagenomic reads from deeply sequenced and less deeply sequenced samples, respectively (Supporting Information Table S2). Three samples (control replicate III on day 1 and HIF-treatment replicate II on days 1 and 3) were removed from further analysis because of their low number of reads (< 10,000 reads). Without quality filtering, all metagenomic reads of each sample (43 samples in total) were individually assembled and scaffolded using SPAdes (Bankevich *et al*., 2012) with default k-mer lengths. Scaffolds longer than 1 kb were used for subsequent analyses. Among these, scaffolds with lengths between 1 kb and 10 kb were used only for abundance estimation. Scaffolds longer than 10 kb were considered partial or almost complete genomes and then used for host prediction, abundance estimation, and functional characterization.

Prokaryotic scaffolds contaminated in the viral fractions were detected using VirSorter2 and then removed (Guo *et al*., 2021). To remove redundancy, these virus-derived scaffolds were clustered into viral operational taxonomic units (vOTUs) based on the average nucleotide identity (ANI) (>95%) using cd-hit (Fu *et al*., 2012). The quality of the vOTUs was assessed using CheckV (Nayfach *et al*., 2021) with default settings. Although low-quality vOTUs are likely to split fragments of viral genomes, leading to overestimation of the number of vOTUs, these were also retained together with high- and middle-quality scaffolds, and we did not use them to compare the number of vOTUs among treatments. The vOTUs judged as “host” regions by CheckV were removed in this step. Consequently, 184 vOTUs generated from the original seawater sample and 296 vOTUs from the microcosm samples were retained (Supporting Information Table S3 and S4). For taxonomic classification of the vOTUs, they were assigned to genus-level genomic OTUs (gOTUs) by ViPTree, as described previously (Nishimura, Watai, *et al*., 2017; Nishimura, Yoshida, *et al*., 2017; Tominaga *et al*., in press).

### Host prediction of vOTUs obtained in this study

The host prediction of these vOTUs was performed following a previously described method (Edwards *et al*., 2016; Tominaga *et al*., 2020, in press). Briefly, putative host groups were assigned based on their similarity to the viral genomic sequence set collected in previous studies (Nishimura, Watai, *et al*., 2017; Tominaga *et al*., in press). If a vOTU was classified into the gOTU, which included host-known viruses, the host group was assigned to the vOTU. Additionally, for vOTUs that were not classified into any host-known gOTUs, host prediction was performed based on nucleotide sequence similarity, CRISPR-spacer matching, and tRNA matching with prokaryotic genomes, as described previously (Tominaga *et al*., 2020, in press).

### Read mapping for abundance estimation of vOTUs

To estimate the relative abundance of each virus, we performed read mapping analysis. To improve the annotation of each read, mapping accuracy, and mapping rate, we used all 79,391 scaffolds that were longer than 1 kb obtained in this study and 10,759 previously published viral genomes (PVG) (8,948 environmental viral genomes obtained from previous studies (Nishimura, Watai, *et al*., 2017; Tominaga *et al*., in press) and 1,811 reference viral genomes (Nishimura, Watai, *et al*., 2017) registered in the NCBI RefSeq database) as reference sequences. Although some of the 79,391 scaffolds were classified into host or viral sequences by VirSorter2 and CheckV, as described above, all the scaffolds including host regions were retained in this step to improve the accuracy of read annotation and read mapping. The reference sequences were clustered into 56,872 of non-redundant genomes based on ANI (>95%) using cd-hit. Before mapping, the quality-control procedure of raw reads was performed using fastp (Chen *et al*., 2018) (reads were scanned with four base pairs of sliding window from the 5’ end and the residual base pairs were dropped if the mean quality score was below 20). These quality-controlled reads were mapped onto the 56,872 scaffolds using Bowtie2 (Langmead and Salzberg, 2012) with a parameter “--score-min L,0,-0.3”. Fragments per kilobase per mapped million reads (FPKM) were calculated using CountMappedReads2 (https://github.com/yosuken/CountMappedReads2). We regarded the reads mapped on scaffolds or genomes which were judged as “viral” by both VirSorter2 and CheckV as virus-derived reads and the reads mapped on the rest of scaffolds as host-derived reads. The reads recruited on neither the host nor the viral scaffold could be regarded as potential virus-derived reads that were not used for assembly.

### Analysis of vOTU dynamics

To analyze the dynamics of the absolute viral density, we estimated the approximate particle number (APN), as described in the prokaryotic community analysis. First, we clustered > 10 kb viral scaffolds including vOTUs obtained in this study and PVGs which were judged as “viral” by both VirSorter2 and CheckV with the 95% identity of ANI again, then applied the formula shown below to the resultant 9,413 vOTUs. 1) We calculated the relative abundance of each of the 9,413 vOTUs. The FPKM value of each vOTU was divided by the total FPKM value of > 1 kb of the viral scaffolds. 2) FPKM-based relative abundance was calibrated using the number of potential virus-derived reads. 3) The calibrated FPKM-based relative abundance was multiplied by the total abundance of viral particles.

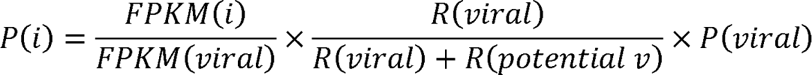

where, *P(i)*: Approximate particle number of vOTUi. *FPKM(i)*: FPKM value of vOTUi. *FPKM(viral)*: total FPKM value of >1 kb viral scaffolds. *R(viral):* the number of reads mapped on > 1kb viral scaffolds. *R(potential v):* number of potential virus-derived reads (reads were not recruited on either host or viral scaffolds). *P(viral):* total abundance of viral particles.

Next, we performed a statistical analysis based on the approximate particle number to detect vOTUs that increased during the microcosm experiments. The approximate particle number of each vOTU in the day 0 samples was compared with that after day 1 by LEfSe (day 0 and after day 1 were set as class and the triplicate flasks were set as subclass). vOTUs that showed a significantly higher density after day 1 were retained. Previous studies have reported that the average burst size of marine bacteria is 50–185 particles/mL (Heldal and Bratbak, 1991; Børsheim, 1993), and an approximate value of their average, 100 particles, was used to determine whether a vOTU increased in terms of approximate particle number. The vOTUs whose approximate particle number did not exceed 100 particles/mL on any day were discarded.

### Gene prediction and annotation of vOTUs

The prediction of open reading frames (ORFs) present in the vOTUs using GeneMarkS (Besemer *et al*., 2001) and homology search of detected ORFs against the NCBI/nr database using GHOSTX (Suzuki *et al*., 2014) were performed using the ViPtree web server (https://www.genome.jp/viptree/). The prediction of AMGs in significantly increased vOTUs was performed using DRAMv (Shaffer *et al*., 2020) with default settings.

### Survey of vOTU abundance in virome data from Osaka Bay natural seawater samples

To confirm whether these retained vOTUs exist in the environment, their abundance in samples collected monthly from May, 2015–November, 2016, in Osaka Bay (Tominaga *et al*., in press) was analyzed by read mapping; a vOTU was regarded as present if even one read was mapped onto it. Raw metagenomic reads were obtained from the DNA Data Bank of Japan (DDBJ) under the project number PRJDB10879 (Tominaga *et al*., in press). Quality control of the reads were performed as described above and the metagenomic reads were mapped to the sequence of vOTUs of interest with 95% identity using Bowtie2 with a parameter “--score-min L,0,-0.3”. The FPKM values were calculated as described above.

### Co-occurrence network analysis between ASVs and vOTUs in Osaka Bay natural seawater samples

Spearman correlations between ASVs and vOTUs of interest were calculated based on their abundance in Osaka Bay samples using the local similarity analysis program (Xia *et al*., 2011). Only samples that covered both prokaryotic and viral datasets were used for this analysis (17 samples), and time delay was not permitted. ASV-vOTU pairs that met r > 0.6 (*p* < 0.01, *q* < 0.05) were regarded as having a significant positive correlation.

### Data availability

The sample information obtained in this study was deposited in the DNA Data Bank of Japan (DDBJ) under project number PRJDB14359. Raw sequence reads can be found under accession number DRA014887.

## Result

### Abundance shifts of prokaryotic cells and viral particles in microcosms

First, we investigated the effects of dissolved intracellular fractions of *Chaetoceros* sp. and *H. akashiwo* (CIF and HIF, respectively) on the growth of prokaryotes in the natural seawater of Osaka Bay. Cell density in all treatments showed similar dynamics and abundance (Fig. 1A). While the cell densities decreased (the control and CIF-treatment) or slightly increased (HIF-treatment) from day 0 to day 1, all the treatments reached their maximum (1.52 ± 0.18 × 10^6^ – 2.78 ± 0.19 × 10^6^ cells/mL) by day 6 followed by stagnation. Based on cell density dynamics, we defined days 0–1 as early when cells were decreasing or slightly increasing, days 2–4 as middle when cells were increasing, and days 5–7 as late when cell densities were almost saturated. While the cell densities of both the HIF- and CIF-treatments were higher than those of the control from the middle phase, only the difference in cell densities between the HIF treatments and the control in the late phase became statistically significant (Mann–Whitney U test: *p* < 0.01). This result suggested that HIF promotes prokaryotic growth, which is consistent with the increase in prokaryotic cells observed in both natural and experimental blooms of phytoplankton in previous studies (Buchan *et al*., 2014; Tada and Suzuki, 2016).

**Figure 1.**
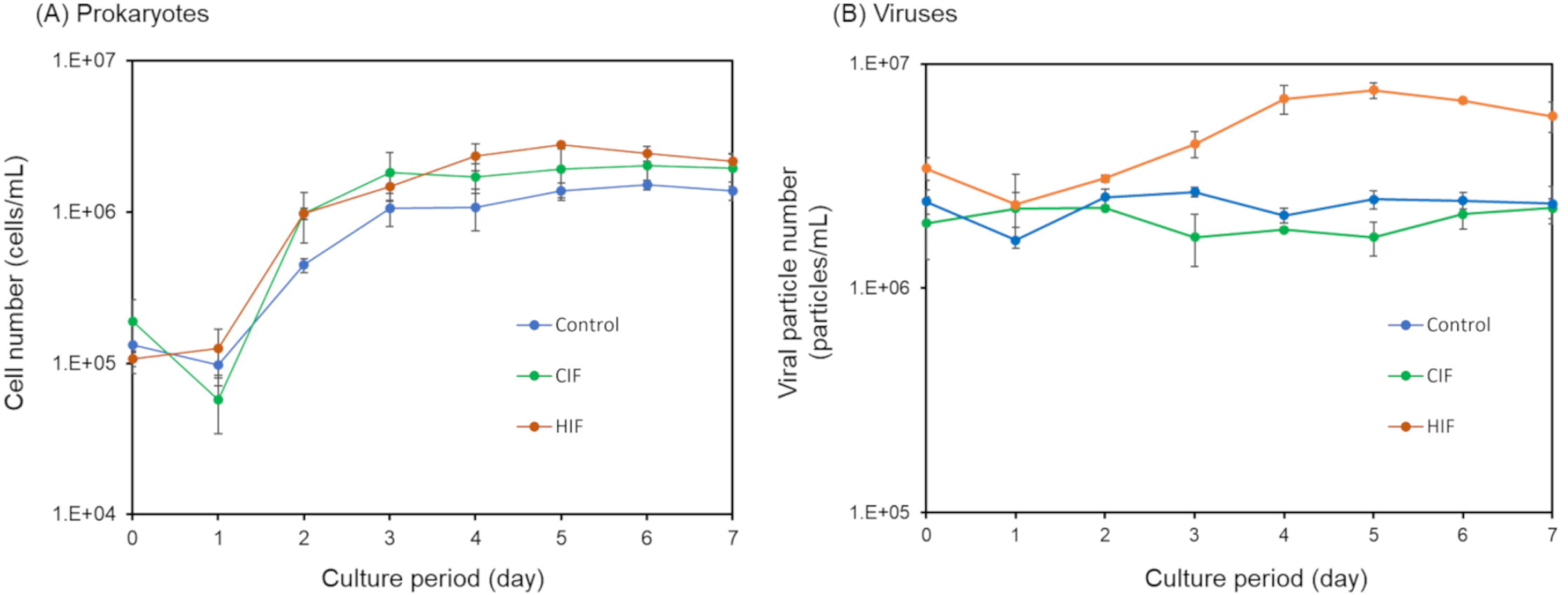
Shifts in abundance of (A) prokaryotic cells and (B) viral particles during the microcosm experiment. Cell and viral counts were obtained using flow cytometry. The average cell and viral numbers in the triplicate flasks are shown. Error bars indicate standard deviation. CIF: *Chaetoceros* sp. intracellular fraction, HIF: *Heterosigma akashiwo* intracellular fraction.

While the viral abundance of control and CIF-treatment samples were stable during the whole culture period (Fig. 1B; 2.34 ± 0.38 × 10^6^ particles/mL in control and 2.01 ± 0.50 × 10^6^ particles/mL in CIF-treatment, on average), that of HIF-treatment samples increased from day 1 to day 5 (2.36 ± 1.47 × 10^6^ to 7.63 ± 1.06 × 10^6^ particles/mL). The viral particle abundance in HIF-treatment samples was significantly greater than that in the control and CIF-treatment samples during the middle and late phases (Mann–Whitney U test: *p* < 0.01). The abundance of these prokaryotic cells and viral particles was used to estimate the absolute dynamics of particular taxa in subsequent analyses.

### Shifts in prokaryotic community structure in response to organic matter

Next, 16S rRNA gene amplicon sequencing analysis confirmed that the dynamics of the marine prokaryotic community were triggered by the intracellular fractions obtained from different phytoplankton species. A total of 415 ASVs were obtained from the original seawater sample. During the microcosm experiments, 1,017 ASVs were obtained in the control and 1,049, and 743 ASVs were obtained on average per flask in CIF- and HIF-treatments, respectively (Supporting Information Table S1). The difference in ASV composition during cultivation was visualized using principal coordinate analysis (PCoA) based on Bray–Curtis dissimilarity (Fig. 2). While all the samples on day 0 were closely plotted with the original seawater sample, the samples in the middle and late phases with the same treatments were plotted closer to each other but separated from those with any other treatment (ANOSIM, *p* < 0.01) and even from their own early phases (ANOSIM, *p* < 0.01). The PCoA indicated that prokaryotic community structures explicitly shifted prior to the middle phase, depending on the presence or absence of the phytoplanktonic dissolved intracellular fraction and the phytoplanktonic species as the source of the fraction.

**Figure 2.**
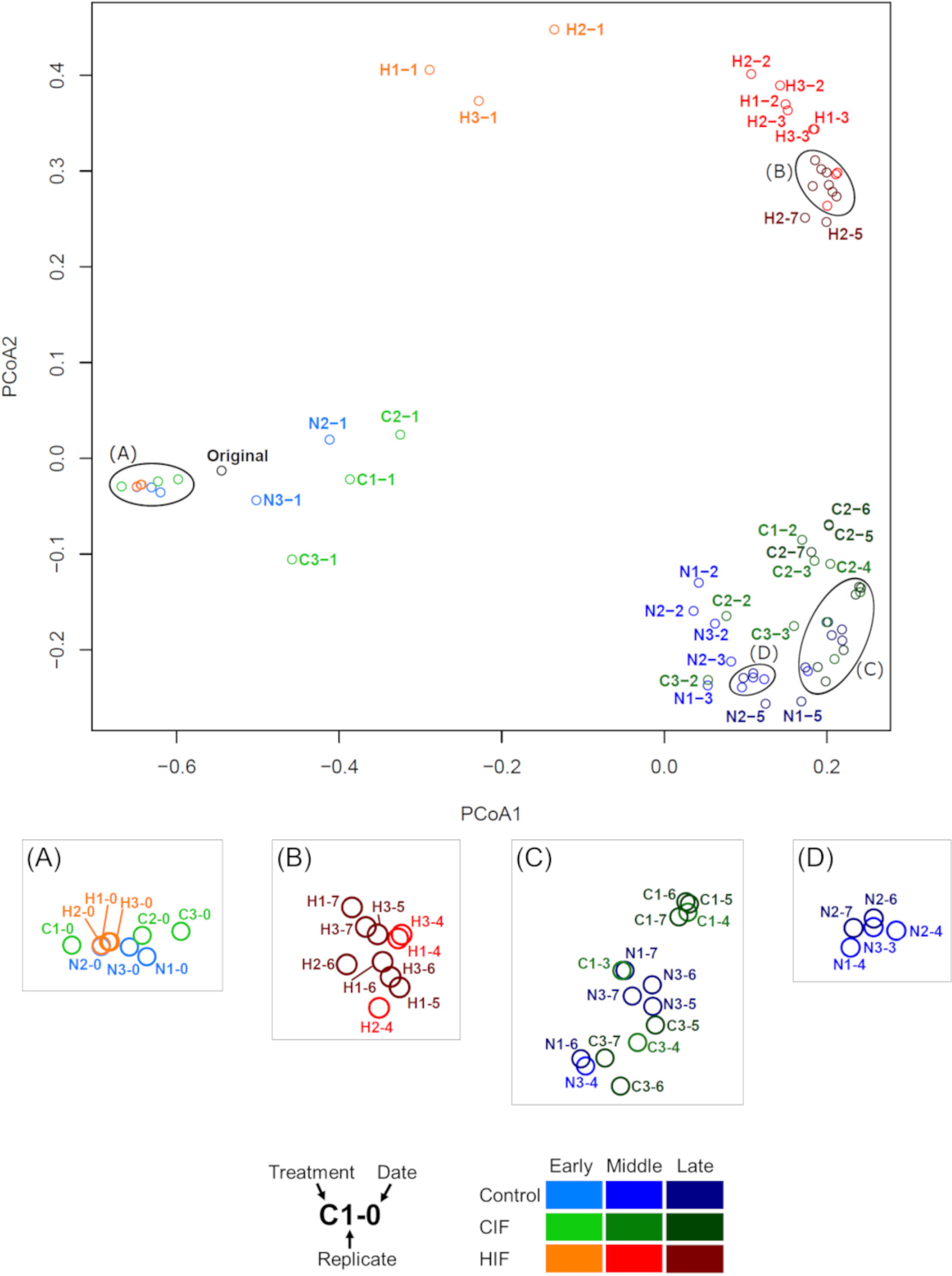
Comparison of ASV compositions among control, CIF-, and HIF-treatments. The number of sequences of each sample were rarified into 9,450 reads prior to the analyses. Bray–Curtis dissimilarity among all samples was illustrated by Principal Coordinate Analysis (PCoA). Samples are distinguished by colors based on the treatments and culture periods. Enlarged figures of areas (A)–(D) are shown separately. ASV: amplicon sequence variants, CIF: *Chaetoceros* sp. intracellular fraction, HIF: *Heterosigma akashiwo* intracellular fraction.

Thus, we analyzed the dynamics of dominant phyla (class level for Proteobacteria) in each treatment (Supporting Information Fig. S2). All the samples on day 0 were dominated by Alphaproteobacteria and Gammaproteobacteria (27.1%–28.8% and 28.8%–34.1%, respectively), followed by Bacteroidetes (18.6%–23.0%). After day 0, these three bacterial groups were still dominant, but their proportions differed among the treatments, supporting the PCoA plotting patterns (Fig. 2).

### Dynamics of abundant ASVs in response to organic matter

We focused on the dynamics of prokaryotes at the ASV level that grew during each treatment. Since increases and decreases in relative abundance during cultivation do not necessarily indicate cell growth and death, we analyzed the dynamics of abundant ASVs by the ACN calculated with the total cell number and relative abundance, as described in the Experimental Procedures section.

We obtained 18, 23, and 23 abundant ASVs in the control, CIF-, and HIF-treatments, respectively. Sixteen of these ASVs were shared between two or three treatments (Supporting Information Fig. S3, Table S5). Among these, 8 ASVs that became abundant in all treatments could be capable of growing independently from phytoplanktonic intracellular fractions (Fig. 3, Supporting Information Fig. S4). Three ASVs that were found to be abundant in both the CIF- and HIF-treatments were regarded as those that were capable of thriving by using both intracellular fractions.

**Figure 3.**
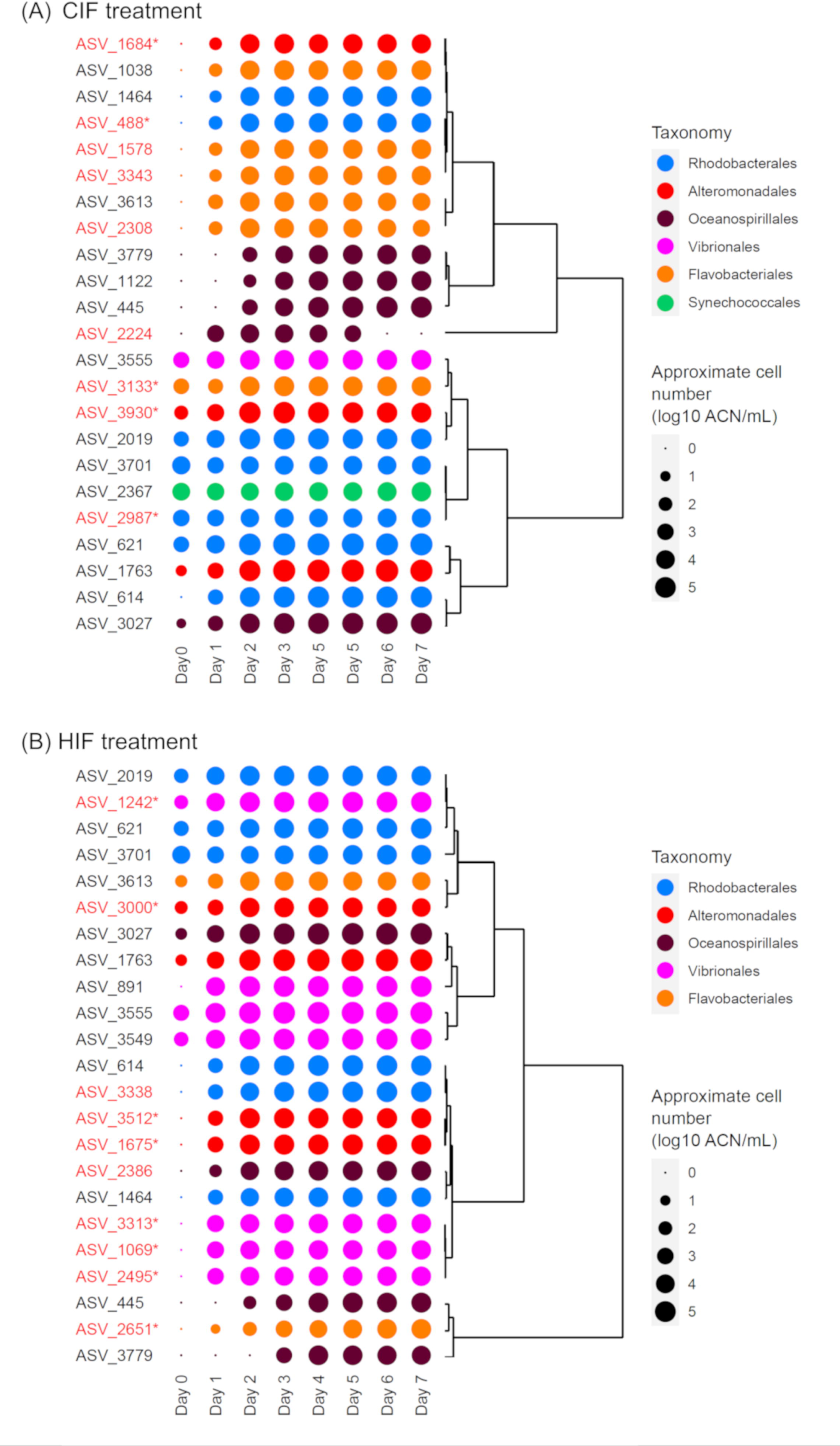
Dynamics of ASVs which were found to be abundant in (A) CIF and (B) HIF samples. Averages of approximate cell number (ACN) in the triplicate flasks are shown as plots in the log scale. The plot colors show order-level taxonomy of each ASV. Dendrograms represent similarity of CAN dynamics among ASVs. ASVs abundant only in the CIF- or HIF-treatments are highlighted in red, and if they are treatment-specific ASVs, indicated by asterisks. ASV: amplicon sequence variants, CIF: *Chaetoceros* sp. intracellular fraction, HIF: *Heterosigma akashiwo* intracellular fraction.

There were nine and 10 ASVs abundant only in either CIF- or HIF-treatments, respectively (Supporting Information Fig. S3, Table S5). To confirm whether these ASVs were specifically abundant in the CIF or HIF-treatments, we performed a differential abundance test using LEfSe. As a result, five and eight ASVs were significantly abundant in the either CIF- or HIF-treatments, respectively. As these were hypothesized to respond to the CIF or HIF, we referred to them as CIF-specific and HIF-specific ASVs, respectively (Table 1). According to ACN, the CIF- and HIF-specific ASVs detected above were suggested to start growing within 1 day from day 0 (Fig. 3), followed by saturation by the end of the middle phase.

**Table 1.** Taxonomic assignment of CIF- and HIF-specific ASVs. Regarding ASVs whose sequence showed 100% identity with cultured bacteria deposited in RefSeq NR database, the hit bacterium is indicated with the corresponding accession number. ASV: amplicon sequence variants, CIF: Chaetoceros sp. intracellular fraction, HIF: Heterosigma akashiwo intracellular fraction.

| ASV ID |  | Taxonomy |  |  | Confidence | Top hit organism<br>(Accession number) |
| --- | --- | --- | --- | --- | --- | --- |
|  |  | Order | Family | Genus |  |  |
| CIF-specific | ASV_1684 | Alteromonadales | Alteromonadaceae |  | 1.000 |  |
|  | ASV_2987 | Rhodobacterales | Rhodobacteraceae | *Planktomarina* | 0.998 | *Planktomarina*<br>*temperata* RCA23<br>(NR_117309,<br>NR_125550) |
|  | ASV_3133 | Flavobacteriales | Flavobacteriaceae | *Polaribacter* | 0.987 |  |
|  | ASV_3930 | Alteromonadales | Flavobacteriaceae | *Glaciecola* | 0.858 | *Glaciecola amylolytica*<br>strain THG-3.7<br>(NR_171501) |
|  | ASV_488 | Rhodobacterales | Rhodobacteraceae | *Nereida* | 0.981 | *Nereida ignava* strain<br>CECT5292<br>(NR_115897)<br>*Nereida ignava* strain<br>2SM4 (NR_042283) |
| HIF-specific | ASV_1069 | Vibrionales | Vibrionaceae | *Vibrio* | 0.837 | *Vibrio splendidus* strain<br>LMG 4042<br>(NR_114888) |
|  | ASV_1242 | Vibrionales | Vibrionaceae | *Vibrio* | 0.763 |  |
|  | ASV_1675 | Alteromonadales | Pseudoalteromonadaceae | *Pseudoalteromonas* | 1.000 |  |
|  | ASV_2495 | Vibrionales | Vibrionaceae | *Vibrio* | 0.982 |  |
|  | ASV_2651 | Flavobacteriales | NS9 marine group | NS9 marine group | 0.920 |  |
|  | ASV_3000 | Alteromonadales | Shewanellaceae | *Psychrobium* | 0.939 |  |
|  | ASV_3313 | Vibrionales | Vibrionaceae | *Vibrio* | 0.978 | *Vibrio gigantis* strain<br>LGP 13 (NR_114910) |
|  | ASV_3512 | Alteromonadales | Pseudoalteromonadaceae | *Pseudoalteromonas* | 0.704 | *Pseudoalteromonas*<br>*marina* strain mano4 |

### Taxonomic characteristics of CIF- and HIF-specific ASVs

The taxonomic compositions of CIF- and HIF-specific ASVs were different at the genus level. The CIF-specific ASVs (Table 1) included *Nereida* (ASV_488, Rohodobacteriales), *Planktomarina* (ASV_2987, Rohodobacteriales), Alteromonadaceae (ASV_1684, genus unassigned), *Glaciecola* (ASV_3930, Alteromonadales), and *Polaribacter* (ASV_3133, Flavobacteriales). ASV_488, ASV_2987, and ASV_3930 showed 100% identity of nucleotide sequences with marine bacteria *Nereida ignava* (NR_115897 and NR_042283), *Planktomarina temperata* (NR_117309 and NR_125550), and *Glaciecola amylolytica* (NR_171501), respectively (Table 1). These species or families are known to be associated with diatom blooms or respond to the organic matter of diatoms (Giebel *et al*., 2011; Tada *et al*., 2012; Teeling *et al*., 2012, 2016; Sarmento *et al*., 2013; Buchan *et al*., 2014; Voget *et al*., 2015).

In contrast, the HIF-specific ASVs were classified into the genera *Psychrobium* (ASV_3000, Alteromonadales), *Pseudoalteromonas* (ASV_1675 and ASV_3512, Alteromonadales), *Vibrio* (ASV_1069, ASV_1242, ASV_2495, and ASV_3313, Vibrionales), and NS9 marine group (ASV_2651, Flavobacteriales) (Table 1). Of these, ASV_3512, ASV_1069, and ASV_3313 had 100% nucleotide sequence identity with *Pseudoalteromonas marina* (NR_042981), *Vibrio splendidus* (NR_114888), and *Vibrio gigantis* (NR_114910), respectively (Table 1); *V. splendidus* causes mortality in mussels and oysters (Balbi *et al*., 2013; Romero *et al*., 2014). *P. marina* and the genus *Vibrio* were appeared to respond to the organic matter of *H. akashiwo* in our previous microcosm experiments (Takebe *et al*., 2020). Although not specific to *H. akashiwo*, the abundance of *Psychrobium* and NS9 marine groups of uncultured Flavobacteriales members is also known to be associated with the dynamics of marine phytoplankton in natural environments (Liu *et al*., 2019; John *et al*., 2022).

The taxonomic composition of the CIF- and HIF-specific ASVs suggests that the dissolved intracellular fractions of different phytoplanktonic species promote bloom-responding bacteria belonging to distinct genera.

### Dynamics of prokaryotic viruses following shifts in prokaryotic communities

Given our finding that CIF and HIF have different effects on the prokaryotic community structures and growth of particular prokaryotic taxa, we investigated the viruses present in the microcosm cultures. To detect viruses that explicitly increased during the experiment in each treatment, we first prepared a virome dataset constituted by 9,313 of non-redundant vOTUs datasets (> 10 kb), of which 465 were newly assembled in this study (Supporting Information Tables S3 and S4) and 8,848 were collected from PVGs generated or collected in previous studies (Nishimura, Watai, *et al*., 2017; Tominaga *et al*., in press). We calculated the approximate particle numbers of these viruses in our microcosm samples. Among these, we regarded vOTUs of the viral particles which significantly increased after day 1 when compared to day 0, as the increased vOTUs were associated with the growth of abundant prokaryotes in our microcosm experiment; the majority of the abundant prokaryotic ASVs increased from day 1 (see above, Fig. 3). As a result, 99, 78, and 42 vOTUs passed this criterion in the control, CIF-, and HIF-treatments, respectively (Supporting Information Fig. S5, Table S6). Fifty-five vOTUs were significantly increased in at least two treatments. Six among these that increased in all treatments could be regarded as increasing independently from the phytoplanktonic intracellular fractions (Fig. 4, Supporting Information Fig. S6). Five vOTUs that increased in both CIF- and HIF- treatments were regarded as vOTUs associated with both phytoplanktonic intracellular fractions. In contrast, 41 and 13 vOTUs significantly increased only in either CIF- or HIF-treatments, respectively (Supporting Information Fig. S5, Table S6). We named these vOTUs CIF- and HIF-specific vOTUs, respectively. Note that the definition of the “specific vOTUs” is different from that of the “specific ASVs” (see above). Given these specific vOTUs, different prokaryotic viruses increased when different phytoplankton-derived intracellular fractions were added.

**Figure 4.**
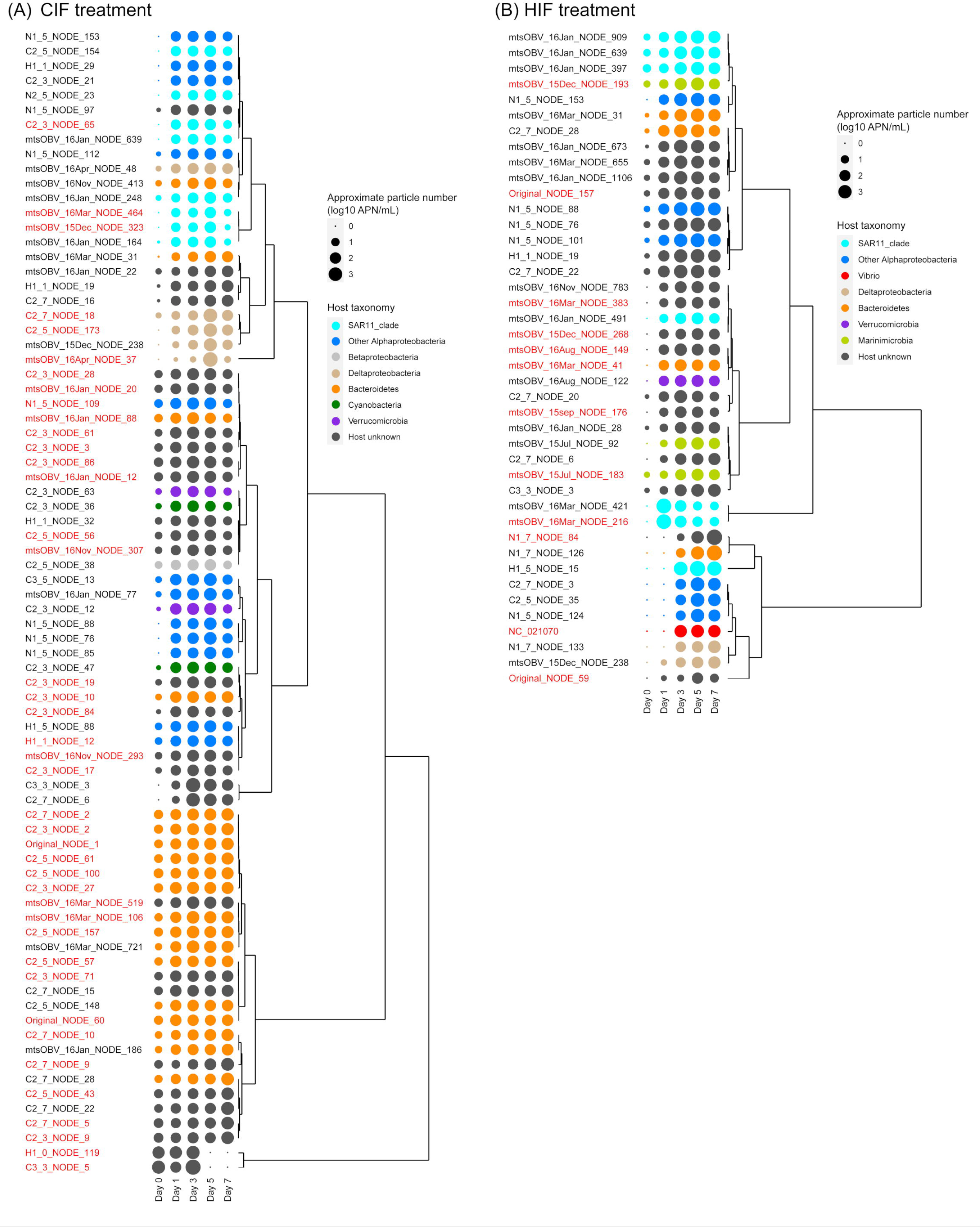
Dynamics of vOTUs which significantly increased in (A) CIF and (B) HIF samples. Averages of approximate particle number (APN) in the triplicate flasks are shown by plots in the log scale. The plot colors show taxonomy of putative host. Dendrograms represent similarity of dynamics of APN among vOTUs. Treatment-specific vOTUs are highlighted in red. vOTU: viral operational taxonomic unit, CIF: *Chaetoceros* sp. intracellular fraction, HIF: *Heterosigma akashiwo* intracellular fraction.

Of the 41 CIF-specific vOTUs, we succeeded in predicting the prokaryotic hosts for 21 (Supporting Information Table S3, S6). Most of them (13 vOTUs; C2_3_NODE_2, C2_3_NODE_10, C2_3_NODE_27, C2_5_NODE_100, C2_5_NODE_157, C2_5_NODE_57, C2_5_NODE_61, C2_7_NODE_10, C2_7_NODE_2, Original_NODE_1, Original_NODE_60, mtsOBV_16Jan_NODE_88, and mtsOBV_16Mar_NODE_106, belonging to g405, g481, g506, g509, and g515, respectively) were predicted as Bacteroidetes virus. The rest belonged to SAR11 clade viruses (3 vOTUs; C2_3_NODE_65, mtsOBV_15Dec_NODE_323, and mtsOBV_16Mar_NODE_464, all belonging to g388), Deltaproteobacteria viruses (3 vOTUs; C2_5_NODE_173, C2_7_NODE_18, and mtsOBV_16Apr_NODE_37, all belonging to g46), and non-SAR11 clade Alphaproteobacteria viruses (H1_1_NODE_12; g389, Rhodobacteriales (Tominaga *et al*., in press) and N1_5_NODE_109; gOTU not assigned, Rhodospirillales (Supporting Information Table S4)) (Supporting Information Table S6).

Five of the gOTUs belonging to the 13 Bacteroidetes vOTUs (g405, g481, g506, g509, and g515) are group 2 Bacteroidetes viruses, one of the most abundant viral groups in the ocean (Tominaga *et al*., 2020). As described above, *Polaribacter* was the only Bacteroidetes (order Flavobacteriales) bacterium that appeared to grow specifically in CIF-treatment (ASV_3133) (Table 1). Notably, the 13 Bacteroidetes viruses did not show sufficiently high similarity in amino acid sequences (Sg < 0.15) with any cultured *Polaribacter* viruses (five strains: NC_028763, NC_028924, NC_062745, NC_062746, and NC_062750, each belonging to g534 or gOTU not annotated). In addition, three of the 13 viruses (C2_3_NODE_2, C2_7_NODE_2, and Original_NODE_1) encoded glycoside hydrolase GH16, a CAZyme that mainly hydrolyzes β-1,3-glucosyl linkages of β-glucan, such as laminarinase (Zverlov *et al*., 2001) (Supporting Information Table S7). This AMG is not present in any genomes of the isolated *Polaribacter* viruses (Kang *et al*., 2015; Bartlau *et al*., 2021) and previously published group 2 Bacteroidetes viruses (Nishimura, Watai, *et al*., 2017; Tominaga *et al*., in press).

Additionally, the CIF-specific ASVs included Rhodobacteriales bacteria, *Nereida* (ASV_488), and *Planktomarina* (ASV_2987) (Fig. 3; Table 1), consistent with the existence of CIF-specific Rhodobacteriales vOTU (H1_1_NODE_12). Although Alteromonadales (Gammaproteobacteria) ASVs were detected as CIF-specific (see above), no CIF-specific vOTU was assigned to an Alteromonadales or Gammaproteobacteria virus (Fig. 4A).

We also succeeded in predicting the hosts for six of the 13 HIF-specific vOTUs (Supporting Information Table S3, S6), including two SAR11 clade viruses (g104; mtsOBV_16Mar_NODE_216 and mtsOBV_16Mar_NODE_421), two Marinimicrobia viruses (g405; mtsOBV_15Dec_NODE_193 and mtsOBV_15Jul_NODE_183), one Gammaproteobacteria virus (g1034; NC_021070), and one Bacteroidetes virus (g504; mtsOBV_16Mar_NODE_41). NC_021070 (Gammaproteobacteria vOTU, g1034) was previously isolated as a virus that infects *Vibrio splendidus* (https://www.ncbi.nlm.nih.gov/nuccore/NC_021070.1/).

*Vibrio* ASVs (ASV_1069, ASV_1242, ASV_2495, and ASV_3313) were detected as HIF-specific (Fig. 3; Table 1), with ASV_1069 being specifically of *V. splendidus.* In addition, the HIF-specific ASVs included ASV_2651 of the NS9 marine group (Bacteroidetes), consistent with the host prediction of the HIF-specific vOTU, mtsOBV_16Mar_NODE_41 (Bacteroidetes vOTU). It is noteworthy that unlike some CIF-specific Bacteroidetes vOTUs, we could not find any AMG in the genomes of the HIF-specific vOTUs. Similar to CIF-treatment, HIF-specific vOTUs assigned to Alteromonadales viruses were not detected, although the HIF-specific ASVs included Alteromonadales ASVs (see above) (Fig. 4B; Supporting Information Table S6).

Although taxonomically different SAR11 vOTUs were detected as CIF- and HIF-specific vOTUs (g388 and g104, respectively) (Fig. 4; Supporting Information Table S6), no SAR11 ASV was identified as abundant ASV. Since four SAR11 ASVs belonging to different clades (clades Ia: three ASVs and clade II: one ASV) were abundant in the original seawater sample (Supporting Information Table S8), the detected SAR11 vOTUs might be derived from an association with any of the SAR11 ASVs. Additionally, although a Rhodospirillales vOTU (N1_5_NODE_109) and Deltaproteobacteria vOTUs (C2_5_NODE_173, C2_7_NODE_18, and mtsOBV_16Apr_NODE_37) were identified as CIF-specific and Marinimicrobia vOTUs (mtsOBV_15Dec_NODE_193, and mtsOBV_15Jul_NODE_183) were detected as HIF-specific (Fig. 4; Supporting Information Table S6), any abundant ASVs both in the original seawater and the microcosm samples were not annotated as the three taxonomic groups of Rhodospirillales, Deltaproteobacteria, and Marinimicrobia (Supporting Information Tables S5 and S8), suggesting that the viruses might also infect prokaryotes of another lineage.

### Co-occurrence of treatment-specific ASVs and vOTUs

Here, we focused on 16 of the treatment-specific vOTUs likely to be infecting Rhodobacteriales, *Vibrio*, and Bacteroidetes, as we detected treatment-specific ASVs in which taxonomic assignments were consistent with the predicted hosts of the above vOTUs in the same treatments (Table 2). We examine consistency between the dynamics of the vOTUs and those of the ASVs in our microcosm samples. Bacteroidetes *Polaribacter* (ASV_3133) and 13 group 2 Bacteroidetes vOTUs (C2_3_NODE_2, C2_3_NODE_10, C2_3_NODE_27, C2_5_NODE_100, C2_5_NODE_157, C2_5_NODE_57, C2_5_NODE_61, C2_7_NODE_10, C2_7_NODE_2, Original_NODE_1, Original_NODE_60, mtsOBV_16Jan_NODE_88, and mtsOBV_16Mar_NODE_106) grew until day 3 (Fig. 3A, 4A). Rhodobacteriales *Nereida* (ASV_488) and *Planktomarina* (ASV_2987) grew until day 2, and Rhodobacteriales vOTUs H1_1_NODE_12 increased from day 0 to day 1. Abundance of the ASVs and the vOTU saturated by the end of the middle phase (Fig. 3A, 4A). *Vibrio* ASVs (ASV_1069, ASV_1242, ASV_2495, and ASV_3313) grew until day 3, followed by saturation. Similarly, *Vibrio* vOTU NC_021070 increased until day 3 (Fig. 3B, 4B). Bacteroidetes NS9, ASV_2651, gradually grew from day 0 to day 7, while mtsOBV_16Mar_NODE_41 increased until day 3, followed by saturation (Fig. 3B, 4B). Collectively, 16 Rhodobacteriales, *Vibrio*, and Bacteroidetes vOTUs showed co-occurrence dynamics with prokaryotic ASVs with the corresponding taxonomy; the vOTUs increased at the same time as or soon after the growth of the prokaryotic ASVs.

**Table 2.**
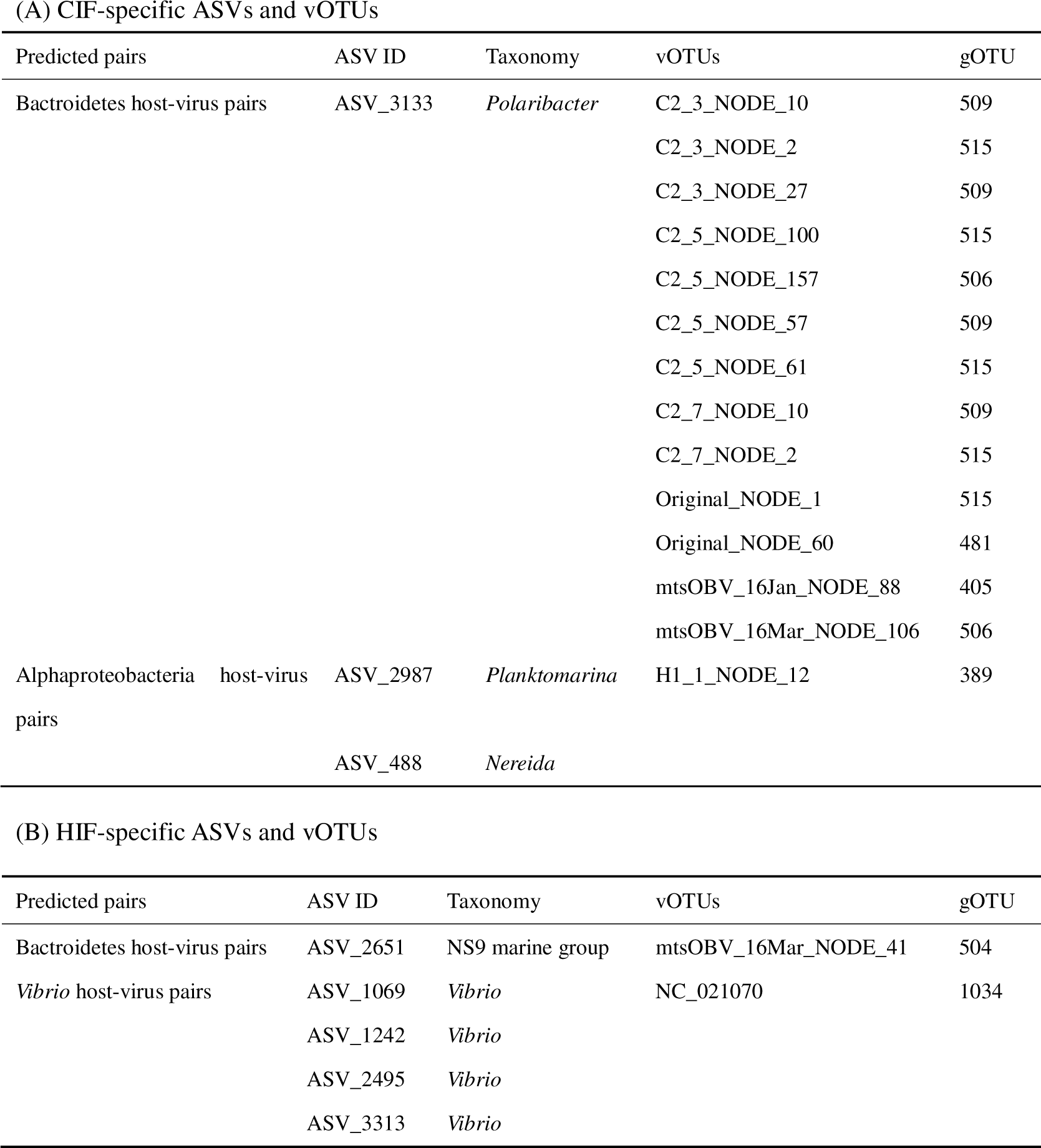
Co-occurence pairs between CIF-specific ASVs-vOTUs and HIF-specific ASVs-vOTUs. ASV: amplicon sequence variants, vOTU: viral operational taxonomic unit, gOTU: genomic operational taxonomic unit, CIF: Chaetoceros sp. intracellular fraction, HIF: Heterosigma akashiwo intracellular fraction. (A) CIF-specific ASVs and vOTUs.

To determine whether the above co-occurrence dynamics of these vOTUs and prokaryotic ASVs could be observed in the environment, we performed a correlation network analysis between them using the 17 datasets from Osaka Bay, collected monthly between 2015–2016 (Tominaga *et al*., in press) (Supporting Information Table S9). We found a significant positive correlation in 13 cases (r > 0.6, *p* < 0.01, *q* < 0.05) (Supporting Information Table S10). ASV_3133 (*Polaribacter*; Bacteroidetes) was detected from May to June, 2015, increased from January to March, 2016, and decreased again by June. Although dynamic patterns were diverse among the Bacteroidetes vOTUs, 11 (C2_3_NODE_2, C2_3_NODE_10, C2_3_NODE_27, C2_5_NODE_157, C2_5_NODE_57, C2_5_NODE_61, C2_7_NODE_10, C2_7_NODE_2, Original_NODE_1, Original_NODE_60, and mtsOBV_16Mar_NODE_106) peaked in May–June, 2015, or January–June, 2016 (Fig. 5A). ASV_2987 (*Planktomarina*) showed similar dynamics to Rhodobacteriales vOTU H1_1_NODE_12, exhibiting an increase in May–June, 2015, and December, 2015–August, 2016 (Fig. 5B). Similarly, *V. splendidus* ASV_1069 and *Vibrio* vOTU NC_021070 increased in January, 2016 (Fig. 5C).

**Figure 5.**
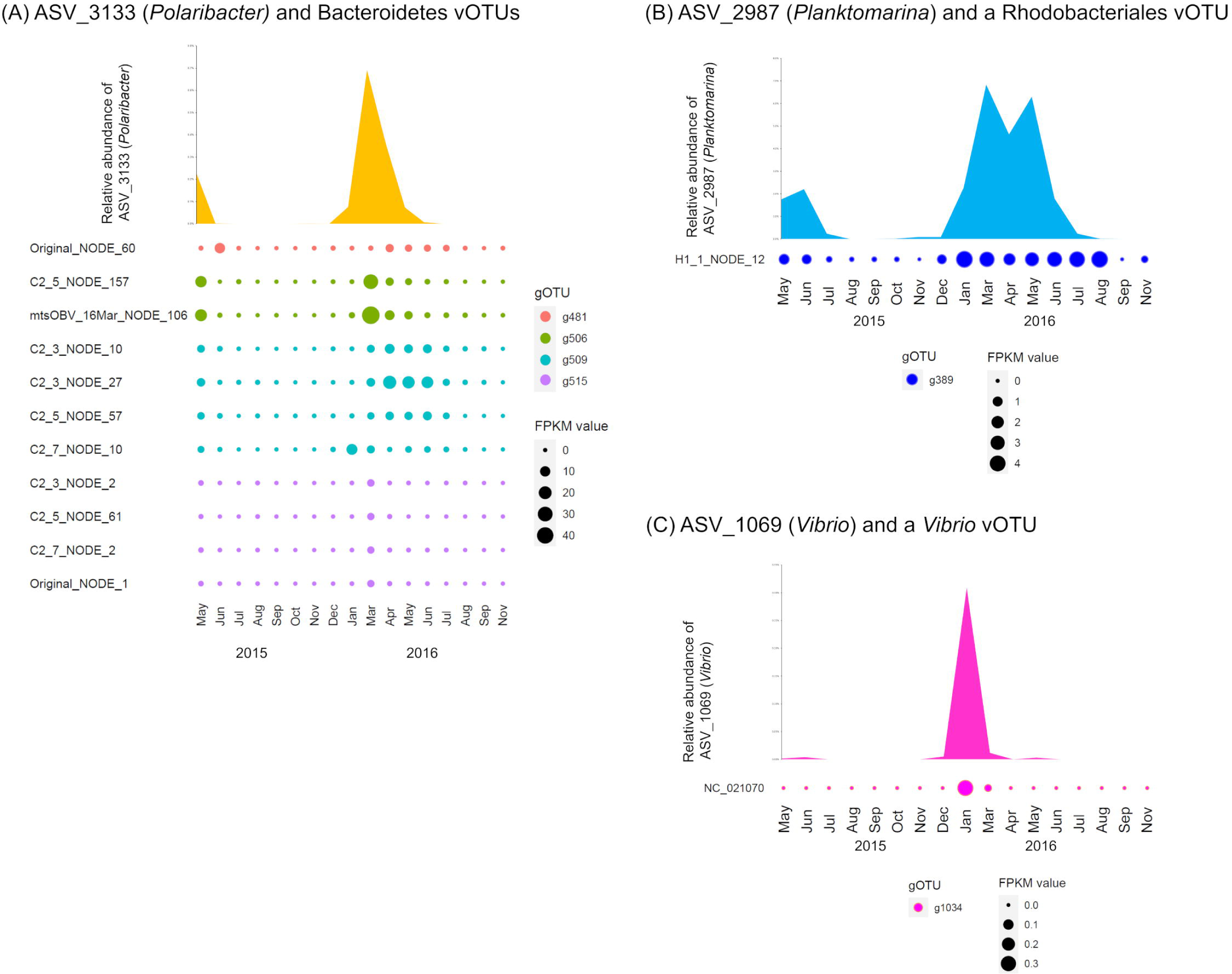
Co-occurrence dynamics of the marine prokaryote viruses and prokaryotes in the Osaka Bay natural seawater samples. (A) ASV_3133 (*Polaribacter*) and Bacteroidetes vOTUs. (B) ASV_2987 (*Planktomarina*) and a Rhodobacteriales vOTU. (C) ASV_1069 (*Vibrio*) and a *Vibrio* vOTU. The environmental datasets monthly collected between May, 2015–November, 2016 (Tominaga *et al.,* in press), were used. Relative abundance of each ASV was calculated by mapping quality-controlled reads of 16S rRNA genes to the ASV sequence with 100% identity using VSEARCH. FPKM value in each vOTU was calculated by mapping quality-controlled viral metagenomic reads to the sequence of the vOTU with 95% identity using Bowtie2. Co-occurrence dynamics are shown only if significantly positive correlation is detected (Spearman correlation; r > 0.6, *p* < 0.01 and Q < 0.05). ASV: amplicon sequence variants, vOTU: viral operational taxonomic unit, CIF: *Chaetoceros* sp. intracellular fraction, HIF: *Heterosigma akashiwo* intracellular fraction, FPKM: fragments per kilobase per mapped million reads.

In addition, as we did not detect Alteromonadales viruses as treatment-specific vOTUs in our microcosm experiments, we subjected the host-unknown, CIF-, and HIF-specific vOTUs (Supporting Information Table S6), and treatment-specific Alteromonadales ASVs (ASV_1684 and ASV_3930 of the CIF-specific ASVs and ASV_1675, ASV_3000, and ASV_3512 of the HIF-specific ASVs) to the same analyses. In the CIF-treatment, 14 host-unknown vOTUs (C2_3_NODE_3, C2_3_NODE_17, C2_3_NODE_19, C2_3_NODE_28, C2_3_NODE_61, C2_3_NODE_71, C2_3_NODE_84, C2_3_NODE_86, C2_5_NODE_56, mtsOBV_16Jan_NODE_12, mtsOBV_16Jan_NODE_20, mtsOBV_16Mar_NODE_519, mtsOBV_16Nov_NODE_293, and mtsOBV_16Nov_NODE_307) started to increase simultaneously with ASV_1684 (genus not annotated) and ASV_3930 (*Glaciecola*) (Fig. 3A, 4A). Similarly, host-unknown HIF-specific vOTUs Original_NODE_59, Original_NODE_157, mtsOBV_15sep_NODE_176, mtsOBV_15Dec_NODE_268, mtsOBV_16Mar_NODE_383, and mtsOBV_16Aug_NODE_149 increased during the growth of ASV_1675 (*Pseudoalteromonas*), ASV_3000 (*Psychrobium*), and ASV_3512 (*Pseudoalteromonas*) (Fig. 3B, 4B). Of these, three of the co-occurrence pairs found in the HIF-treatment showed significant correlative dynamics (r > 0.6, *p* < 0.01, *q* < 0.05) in the natural seawater samples (Supporting Information Table S10): ASV_1675 (*Pseudoalteromonas*), ASV_3512 (*Pseudoalteromonas*)-Original_NODE_59, and ASV_3000 (*Psychrobium*)- mtsOBV_16Mar_NODE_383 (Supporting Information Fig S7). The eight genes in the Original_NODE_59 genome, which have functionally annotated homologs in cellular organisms or viruses, were all similar to the genes present in Alphaproteobacteria virus genomes (GHOSTX; e-value < 1E-8) (Supporting Information Table S11). For instance, the amino acid sequences of proteins encoded in gene1 and gene10 of this vOTU were highly similar to the putative tail fiber protein of *Pelagibacter* phage HTVC033P (MZ892993_66; e-value=1.4E-44) and the putative terminase small subunit of *Roseobacter* phage CRP-212 (MZ892988_42; e-value=3.9E-27). In contrast, genes of mtsOBV_16Mar_NODE_383 had homologs in taxonomically diverse bacteria and viruses (GHOSTX; evalue < 1E-8) (Supporting Information Table S11), and the amino acid sequences of proteins encoded in gene5 and gene10 were similar to those of PD-(D/E)XK nuclease-like domain-containing protein of *Mycobacterium heckeshornense* (WP_071700202, evalue=6.90E-11) (Actinobacteria) and DNA cytosine methyltransferase of *Margalitia camelliae* (WP_101355680, evalue=3.8E-51) (Bacillota), respectively. Amino acid sequences of proteins encoded by gene21 and gene25 of mtsOBV_16Mar_NODE_383 were similar to those of a phage terminase large subunit of *Zunongwangia* phage (WP_084841783, e-value=1.5E-44) and a phage tail tape measure protein of *Lutibacter* phage (WP_144895146, e-value=3.6E-27), both of which are viruses infecting the class Bacteroidia (phylum Bacteroidetes).

## Discussion

In the present study, we combined 16S rRNA gene amplicon and viral metagenomic analyses in our microcosm experiment and demonstrated two important aspects of microbial ecosystems in marine environments. Specific prokaryotic viruses can increase following changes in the prokaryotic community within a few days and their composition was affected by the bloom-forming phytoplanktonic species. We confirmed that the addition of dissolved intracellular fractions derived from different phytoplanktonic species promotes the growth of taxonomically distinct prokaryotes that are known to react to phytoplanktonic organic matter or are associated with blooms. The differentially grown prokaryotes in different phytoplankton-derived intracellular fractions (Fig. 3; Table 1) had an associated increase in distinct prokaryotic viruses (Fig. 4). As the treatment-specific viruses were rarely detected on day 0 in this study (Fig. 4), these viruses were most likely increased by infecting the abundant prokaryotes; if the Kill-the-Winner hypothesis stands, prokaryotic taxa with a faster growth rate are more susceptible to viral infection (Thingstad, 2000). Thus, we succeeded in identifying both known and new possible host-virus pairs in our microcosm experiment (Table 2).

The most probable host-virus pair was *V. splendidus* (ASV_1069) and its virus NC_021070 (https://www.ncbi.nlm.nih.gov/nuccore/NC_021070.1/), which appeared in the HIF-treatment (Fig. 3, 4; Table 2). Preference of *Vibrio* spp. for the intracellular component of *H. akshiwo* was in good agreement with our previous study (Takebe *et al*., 2020). The co-occurrence between them in the Osaka Bay samples further supports the *H. akashiwo*-*V. splendidus*-NC_021070 virus relationship (Fig. 5C; Supporting Information Table S10).

The host-virus relationship between *Polaribacter* ASV_3133 and 13 of group 2 Bacteroidetes viruses in the CIF-treatment were the most prominent examples of possible host-virus pairs (Fig. 3, 4; Table 2), as previously unknown but possible roles of viruses for their host metabolism were elucidated. *Polaribacter* ASV_3133 grew specifically in the microcosm, including in the CIF (Table 1). This is consistent with a previous study that detected *Polaribacter* in natural diatom blooms and their well-known function of utilizing phytoplankton-derived polysaccharides, such as laminarin included in diatom cells, as growth substrates (Teeling *et al*., 2012, 2016; Avcı *et al*., 2020). Importantly, *Polaribacter* ASV_3133 was the only Bacteroidetes ASV detected as CIF-specific, and the 13 group 2 Bacteroidetes viruses were also CIF-specific (Fig. 4; Table 2). Co-occurrence between ASV_3133 and all 13 CIF-specific vOTUs (group 2 Bacteroidetes viruses), except for C2_5_NODE_100 and mtsOBV_16Jan_NODE_88, detected in Osaka Bay samples (Fig. 5A; Supporting Information Table S10) further emphasizes the confidence of these CIF-specific host-virus pairs. Three of these Bacteroidetes vOTUs (C2_3_NODE_2, C2_7_NODE_2, and Original_NODE_1) possessed possible laminarinase (GH16) as the AMG. Considering that diatom cells contain laminarin as the major carbon storage material (Becker *et al*., 2020), these viruses are presumed to obtain the energy required for proliferation by promoting the utilization of sugars generated by laminarin degradation by the host. In addition to the other vOTUs, previously isolated *Polaribacter* viruses derived from a diatom bloom sample (Kang *et al*., 2015; Bartlau *et al*., 2021) lack genes for GH16, suggesting diverse propagation strategies in viruses infecting Bacteroidetes hosts responding to diatom blooms.

Rhodobacteriales (Alphaproteobacteria) bacteria, *Nereida ignava*, and *Planktomarina temperata* are ecologically important because they are abundant, potentially bloom-responding taxa in coastal areas (Giebel *et al*., 2011; Sarmento *et al*., 2013; Voget *et al*., 2015). Although neither *Nereida* nor *Planktomarina* viruses have been isolated (as of Oct 2022 in Virus-Host DB) (https://www.genome.jp/virushostdb/), we identified one potential virus (H1_1_NODE_12) that might be capable of infecting them or either of them. *N. ignava* and *P. temperata* were the only Rhodobacteriales species that were identified as CIF-specific, and H1_1_NODE_12 was predicted to be the CIF-specific Rhodobacterial virus (Table 2). In particular, the co-occurrence between *P. temperata* ASV_2987 and H1_1_NODE_12 in Osaka Bay samples further supports the *P. temperata*-H1_1_NODE_12 virus pair (Fig. 5B; Supporting Information Table S10). These biological interactions among *H. akashiwo,* prokaryotes, and prokaryotic possibly occur in Osaka Bay, where *H. akashiwo* forms blooms.

Our experiment might also capture potential host-virus pairs unknown by previous culture-dependent viral isolation and even by genomics-based *in silico* prediction. The only Bacteroidetes bacterial ASV detected as HIF-specific is of the NS9 marine group (ASV_2651) (Fig. 3; Table 1), which is an uncultured bacterial clade of Bacteroidetes but is abundant in coastal areas (Seo *et al*., 2017; Liu *et al*., 2019). HIF-specific Bacteroidetes vOTU (mtsOBV_16Mar_NODE_41) co-occurred with ASV_2651 (Fig. 4; Supporting Information Table S6) in the microcosm samples, suggesting that they are potential a host-virus pair.

Although Alteromonadales (Gammaproteobacteria) ASVs were detected as CIF-specific and HIF-specific ASVs (Fig. 3), no CIF-specific or HIF-specific vOTU were assigned as Alteromonadales viruses (Fig. 4), possibly due to insufficient viral isolates infecting them and the limitations of current host prediction methods. Indeed, viruses infecting *Glaciecola amylolytica* (ASV_3930, CIF-specific) and *Psychrobium* (ASV_3000, HIF-specific) have not been isolated (as of Oct 2022 in Virus-Host DB) (https://www.genome.jp/virushostdb/). However, host-unknown vOTUs (Original_NODE_59 and mtsOBV_16Mar_NODE_383) in HIF-treatments were correlated with the HIF-specific prokaryote ASVs (*Pseudoalteromonas* ASV_1675, *Psychrobium* ASV_3000, and *Pseudoalteromonas* ASV_3512) in both the microcosm and natural seawater samples (Fig. 3, 4; Supporting Information Fig S7, Table S10), indicating the possibility that the host-unknown CIF- and HIF-specific vOTUs include Alteromonadales viruses. Such a correlation is less confident in predicting host-virus relationships compared to genome-based prediction methods (Edwards *et al*., 2016).

Collectively, based on the difference in predicted host-virus pairs between the treatments, we concluded that globally distributed bloom-forming phytoplanktonic species, *H. akashiwo* and *Chaetoceros*, promote, at least partially, distinct prokaryotic viruses through forming the different composition of bloom-responding prokaryote species. Thus, we propose that the succession of bloom-forming phytoplankton can change the composition of the abundant prokaryotes, resulting in the transition of abundant prokaryotic viruses. Changes in viral composition depending on bloom-forming species, in turn, would affect the dynamics and metabolism of prokaryotes (*e.g.* GH16), altering biogeochemical cycling during blooms. Since our focal phytoplankton, *H. akashiwo* and *Chaetoceros,* were reported to emerge simultaneously or successively in the same coastal area (Needham and Fuhrman, 2016), the prokaryotic responses and viral dynamics observed in this study would further help to understand coastal microbial ecosystems.

## Acknowledgements

Computational analyses were partly performed at the Super Computer System, Institute for Chemical Research, Kyoto University. This study was supported by Grants-in-Aid for Scientific Research (No. 16H06429, No. 17H03850, No. 21H05057, and No. 21J14854) from the Japan Society for the Promotion of Science (JSPS).

