## Supplemental Figures for "Taxonomic difference in marine bloom-forming phytoplanktonic species affects dynamics of both bloom-responding prokaryotes and prokaryotic viruses"

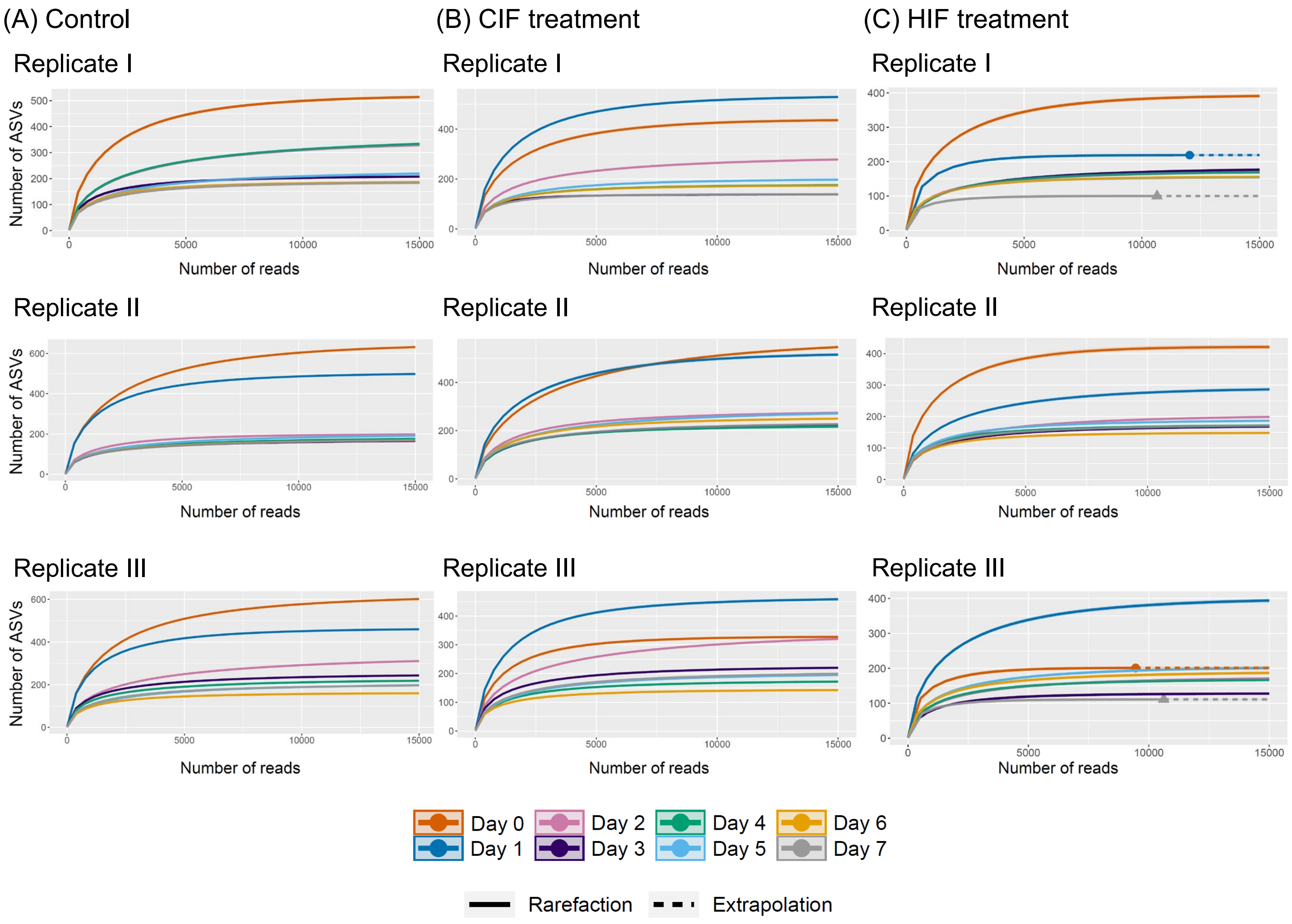


**Supporting Information Figure S1.** Rarefaction curves generated from the number of ASVs detected in each sample**.** (A) **C**ontrol, (B) CIF treatment, and (C) HIF treatment. From each sample, 15,000 reads were randomly extracted. Samples with fewer than 15,000 reads were compensated using the "extrapolation" function in "iNEXT". Samples are distinguished by different colors based on the date of collection. ASV: amplicon sequence variants, CIF: *Chaetoceros* sp. intracellular fraction, HIF: *Heterosigma akashiwo* intracellular fraction.

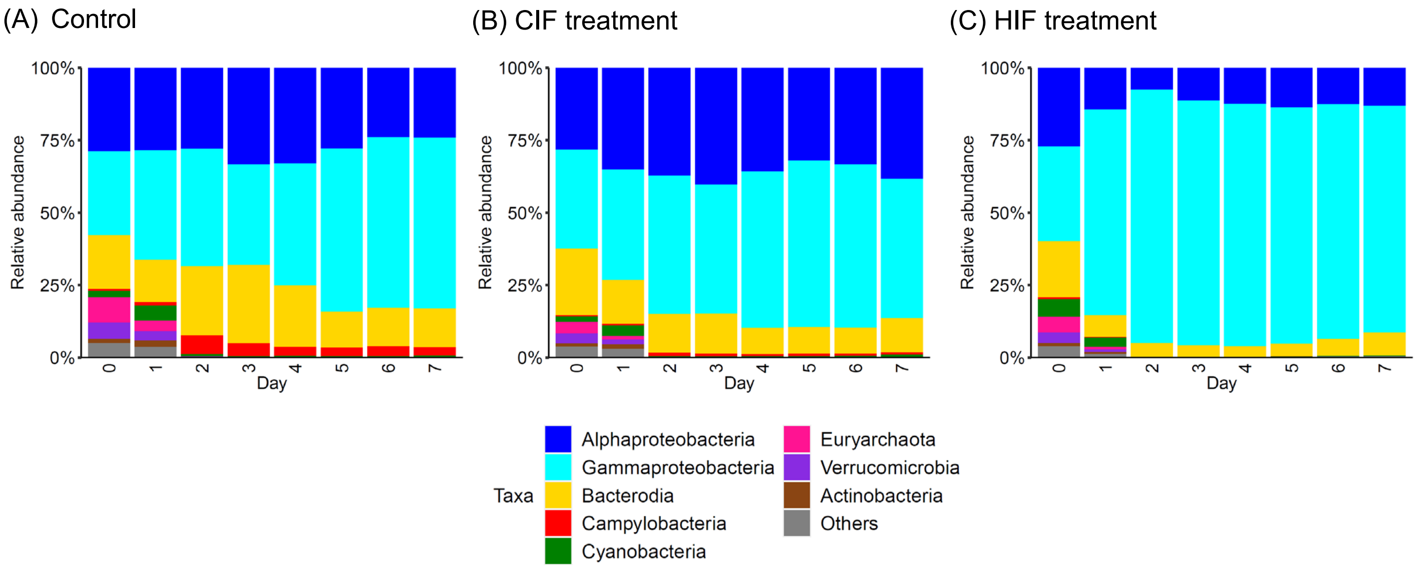


**Supporting Information Figure S2.** Relative abundance of phylum-level (class-level for proteobacteria) phylogenetic groups in the microcosm samples. For each treatment, averaged relative abundance in the triplicate flasks is shown. CIF: *Chaetoceros* sp. intracellular fraction, HIF: *Heterosigma akashiwo* intracellular fraction.

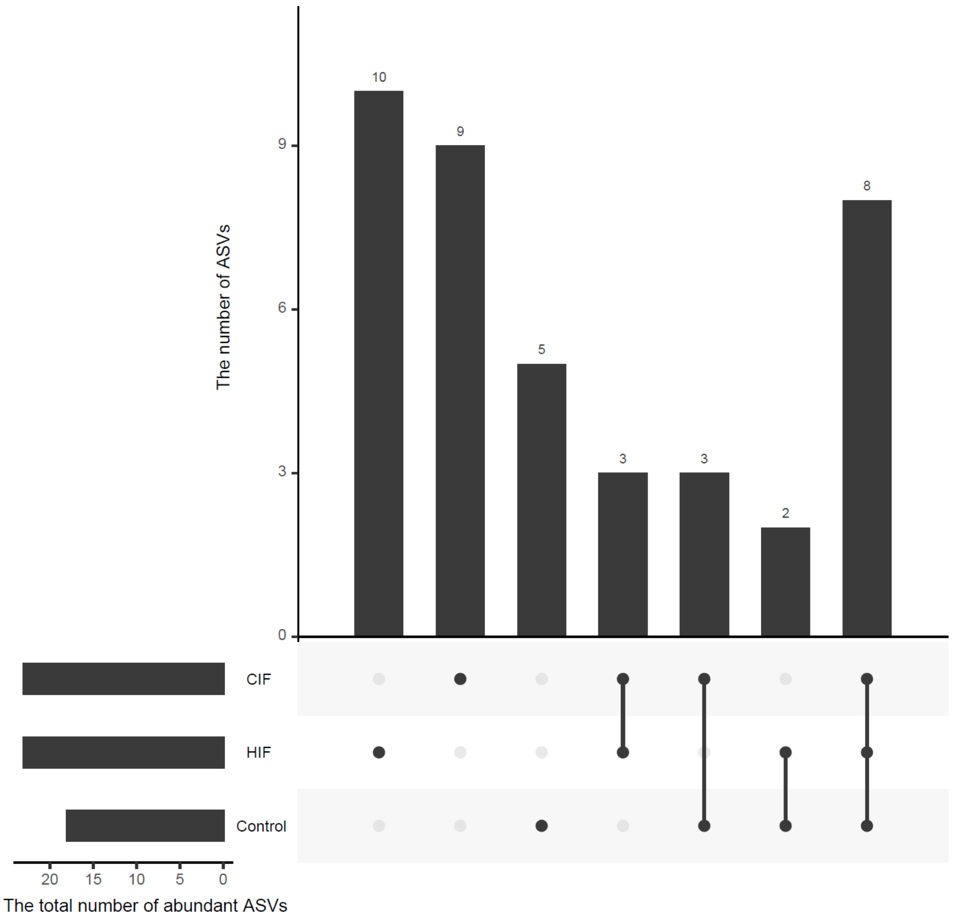


**Supporting Information Figure S3.** UpSet plot indicating the distribution pattern of all the abundant ASVs. The number shown above bar graph indicates those of abundant ASVs detected in each treatment. ASV: amplicon sequence variants, CIF: *Chaetoceros* sp. intracellular fraction, HIF: *Heterosigma akashiwo* intracellular fraction.

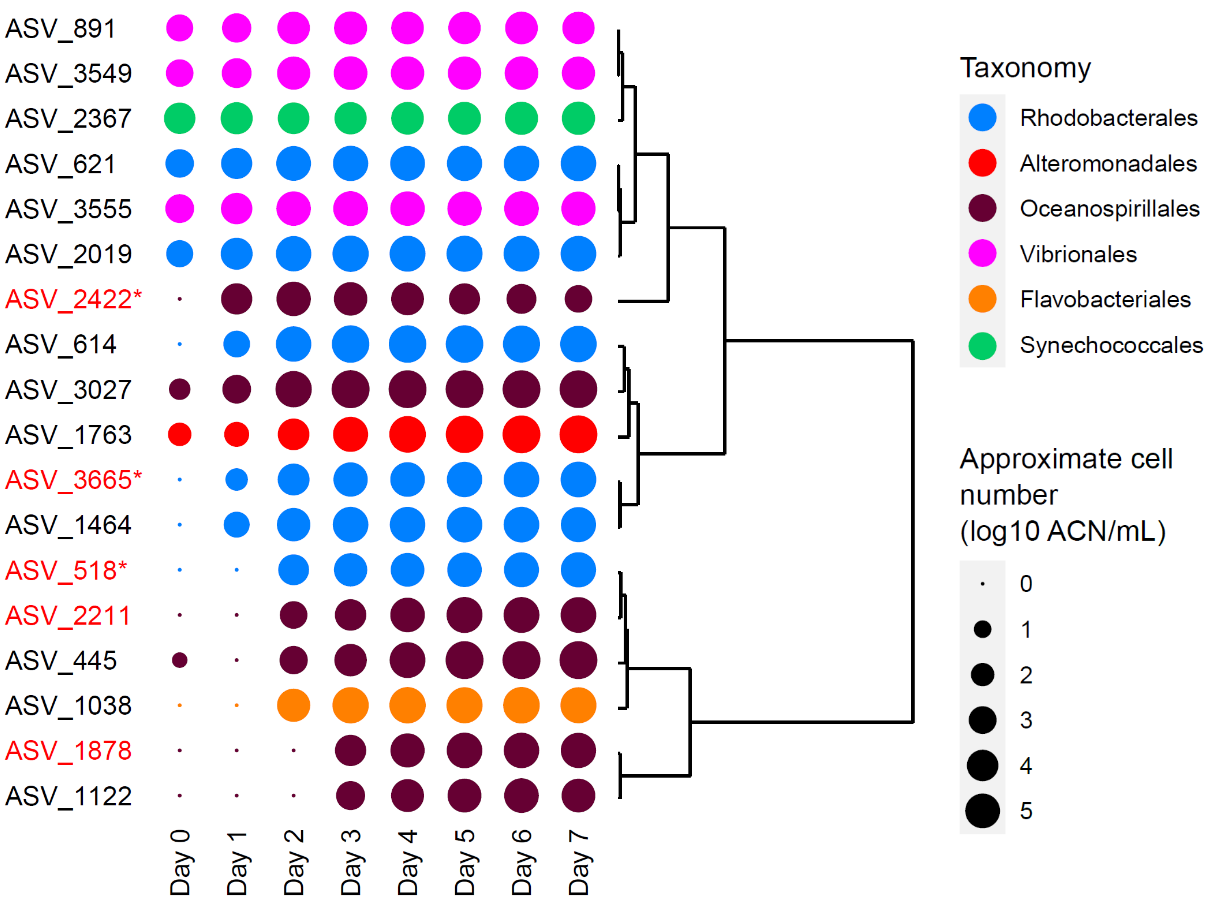


**Supporting Information Figure S4.** Dynamics of ASVs which were found to be abundant in control samples. Averages of approximate cell number (ACN) in the triplicate flasks are shown as plots in the log scale. The plot colors show order-level taxonomy of each ASV. Dendrograms represent similarity of dynamics of ACN among ASVs. ASVs abundant only in the control samples are highlighted in red, and if they are control-specific ASVs, indicated by asterisks. ASV: amplicon sequence variants.

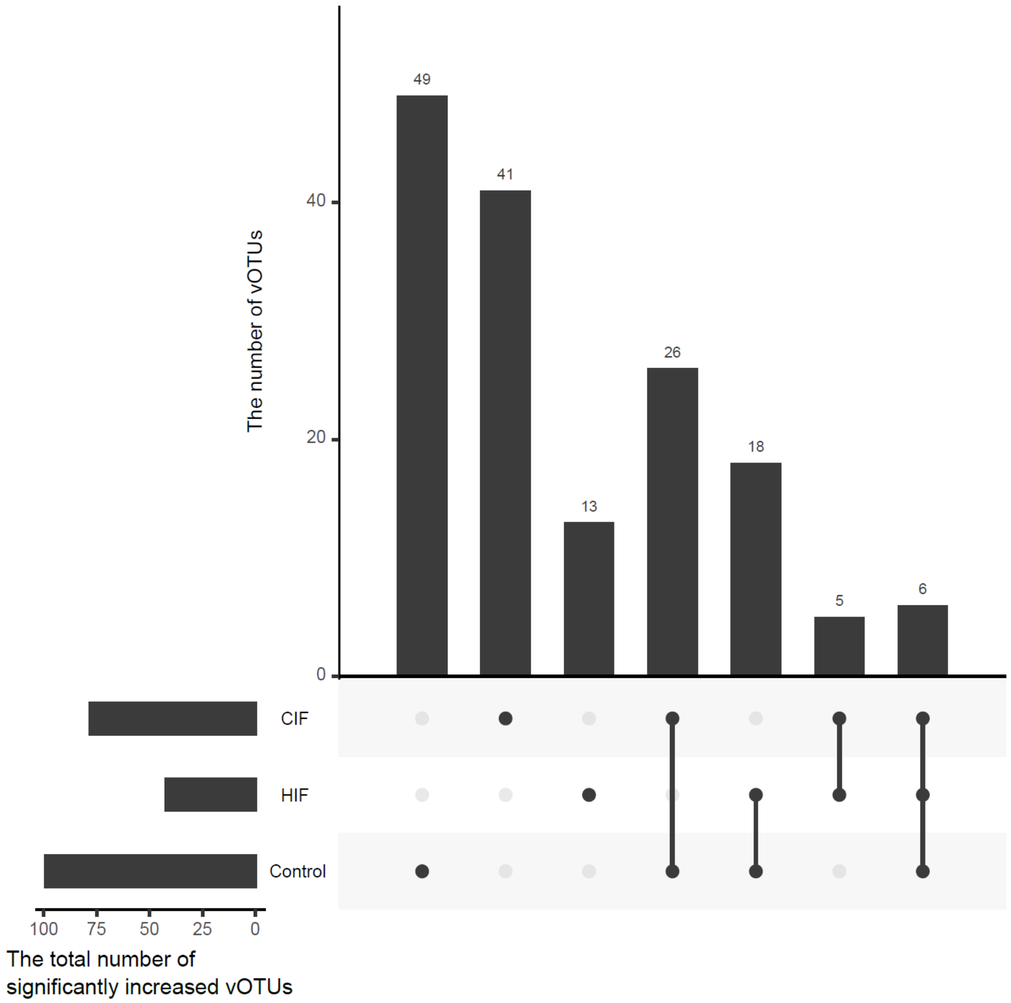


**Supporting Information Figure S5.** UpSet plot indicating the distribution pattern of the significantly increased vOTUs. The number shown above bar graph indicates those of increased vOTUs detected in each treatment. vOTU: viral operational taxonomic unit, CIF: *Chaetoceros* sp. intracellular fraction, HIF: *Heterosigma akashiwo* intracellular fraction.

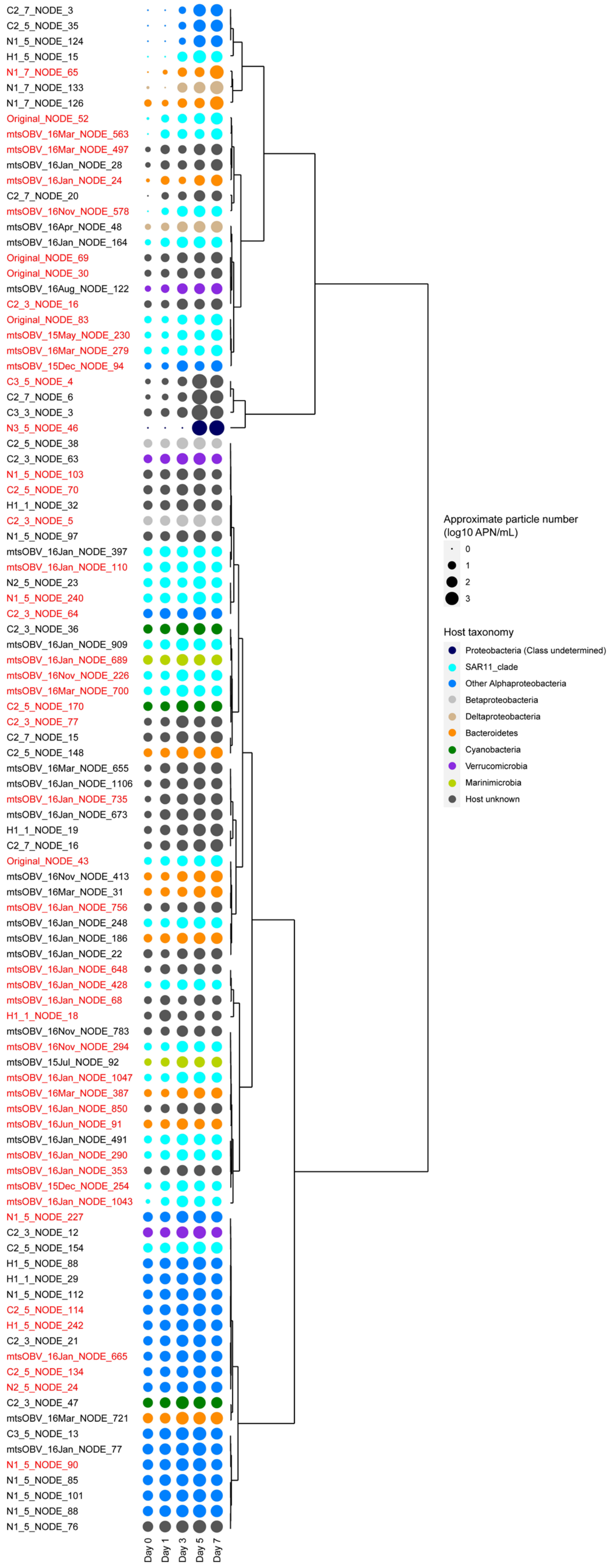


**Supporting Information Figure S6.** Dynamics of vOTUs which significantly increased in the control samples. Averages of approximate particle number (APN) in the triplicate flasks were shown by plots in the log scale. The plot colors show taxonomy of putative host. Dendrograms represent similarity of dynamics of APN among vOTUs. Control-specific vOTUs are highlighted in red. vOTU: viral operational taxonomic unit.

(A) ASV_1675 (*Pseudoalteromonas*) and a host-unknown vOTU

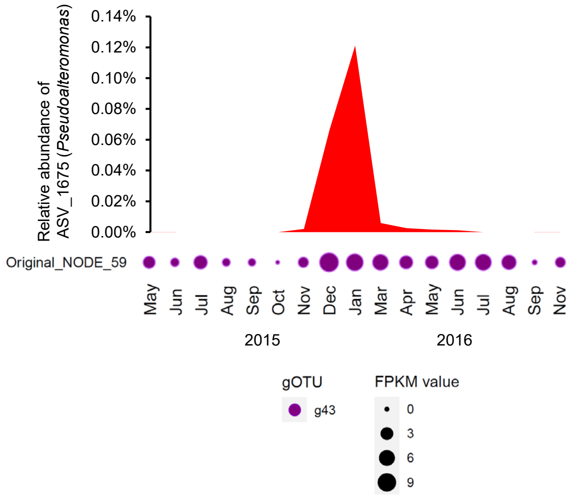


(B) ASV_3000 (*Psychrobium*) and a host-unknown vOTU

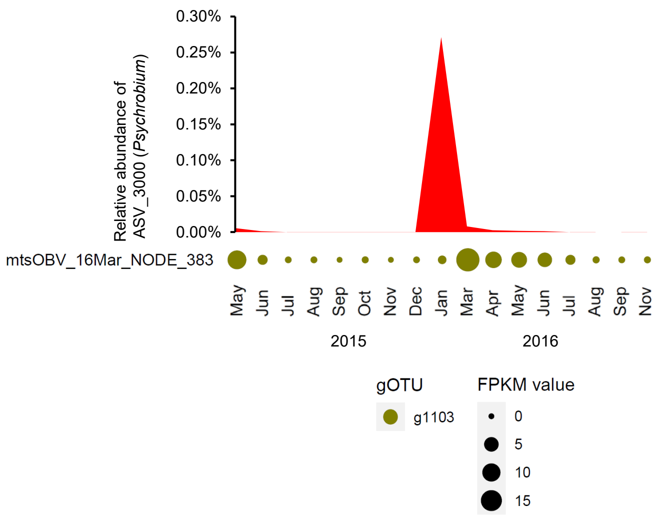


(C) ASV_3512 (*Pseudoalteromonas*) and a host-unknown vOTU

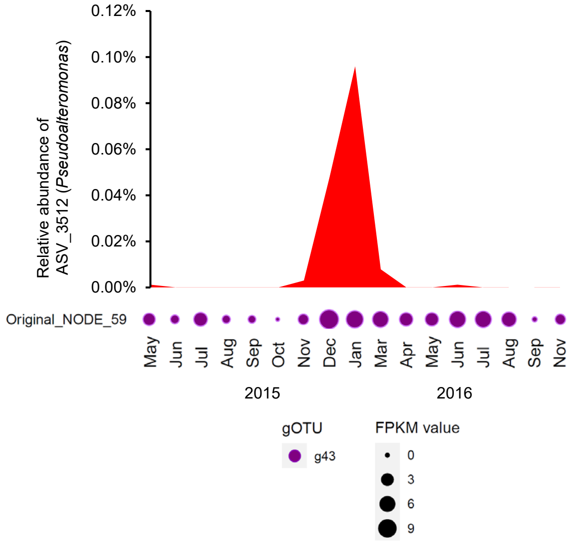


**Supporting Information Figure S7.** Co-occurrence dynamics of the marine prokaryote viruses and the prokaryotes in the Osaka Bay natural seawater samples. (A) ASV_1675 (*Pseudoalteromonas*) and a host-unknown vOTU. (B) ASV_3000 (*Psychrobium*) and a host-unknown vOTU. (C) ASV_3512 (*Pseudoalteromonas*) and a host-unknown vOTU. The environmental datasets monthly collected between May, 2015–November, 2016 (Tominaga *et al.*, in press), were used. Relative abundance of each ASV was calculated by mapping quality-controlled reads of 16S rRNA genes to the ASV sequence with 100% identity using VSEARCH. FPKM value in each vOTU was calculated by mapping quality-controlled viral metagenomic reads to the sequence of the vOTU with 95% identity using Bowtie2. Co-occurrence dynamics are shown only if significantly positive correlation is detected (Spearman correlation; $r > 0.6$, $p<0.01$ and $q<0.05$). ASV: amplicon sequence variants, vOTU: viral operational taxonomic unit, gOTU: genomic operational taxonomic unit, FPKM: fragments per kilobase per mapped million reads.
